# Fusion Oncoprotein EWSR1::FLI1 Invades Nucleosomes at Consensus ETS Motifs and GGAA Microsatellites

**DOI:** 10.64898/2026.09.16.752125

**Authors:** Ruo-Wen Chen, Runwei Zhou, E. John Tokarsky, Megann A. Boone, Andrea K. Byrum, Michael G. Poirier, Emily R. Theisen

## Abstract

Ewing sarcoma is an aggressive bone malignancy occurring in children, adolescents, and young adults. Most cases are caused by expression of the fusion oncoprotein EWSR1::FLI1, which contains the low complexity domain (LCD) of EWSR1 and the DNA-binding domain (DBD) of FLI1. Previous genomic studies indicate EWSR1::FLI1 accesses GGAA microsatellites in chromatin to function as a potent transcriptional regulator. Due to the technical challenges of purifying full-length EWSR1::FLI1, mechanistic studies biochemically characterizing its pioneer activities have been lacking. Here, we purified both full-length EWSR1::FLI1 and truncated DBD constructs to conduct biochemical and fluorescence-based experiments investigating interactions with different motifs in free DNA and nucleosomes. Both truncated and full-length EWSR1::FLI1 show efficient target binding in nucleosomes, and that the fourth alpha-helix in the FLI1 DBD enhances nucleosome-binding efficiency. Surprisingly, we also observe differences in both free DNA binding affinity and sequence preference between truncated and full-length proteins, though these changes are not apparent in nucleosome-binding assays. These findings reveal that EWSR1::FLI1 possesses a key pioneer factor property, efficiently targeting its binding site within nucleosomes, and that full-length EWSR1::FLI1 binding shifts to preferentially target GGAA repeats, even on motifs that bind a single EWSR1::FLI1 protein.

## Introduction

Ewing sarcoma is the second most common bone-associated malignancy occurring in children, adolescents, and young adults (1). Molecularly, Ewing sarcoma is characterized by a pathognomonic chromosomal translocation that fuses a 5’ portion of a FET family gene (*FUS*, *EWSR1*, and *TAF15*) with the 3’ portion of a gene encoding an E26 transformation specific (ETS) transcription factor (TF) from the PEA3 or ERG subfamily. The most common translocation is t(11; 22) (q24; q12), resulting in the formation of the *EWSR1::FLI1* fusion oncogene in 85%-90% of cases (2, 3). The encoded oncoprotein fuses the low complexity domain (LCD) of EWSR1 to the winged helix-turn-helix (wHTH) DNA-binding domain (DBD) of FLI1. The EWSR1 LCD serves as a potent transactivation domain, while the FLI1 DBD confers sequence-specific binding to the EWSR1::FLI1 fusion oncoprotein (3–6). Together, EWSR1::FLI1 functions as an aberrant TF that disrupts normal gene regulation and alters cell fate to promote Ewing sarcoma oncogenesis.

Since EWSR1::FLI1 functions as a master regulator of cell fate, it is proposed to be a pioneer transcription factor (PF) (7). PFs regulate lineage-specific genes and determine cell identity and differentiation status through their ability to recognize and bind their motifs in closed chromatin, facilitate chromatin opening, and promote ATP-dependent chromatin remodeling to enable *cis*-regulatory element activation (8, 9). PFs are characterized by their ability to bind to nucleosomal motifs at affinities within an order of magnitude of those for naked DNA (10–12). In contrast, the binding affinity of canonical TFs can be impeded by nucleosomes over 1000-fold (13, 14).

EWSR1::FLI1 has been shown to bind at both the FLI1 consensus high affinity (HA) motif (5’-*ACCGGAAGTG*-3’) and repetitive *GGAA* microsatellites (15–17). *GGAA* microsatellites are critical EWSR1::FLI1 response elements in both reporter assays and endogenous loci in Ewing sarcoma patients (17–19). EWSR1::FLI1 binding at *GGAA* repeats is a fundamental molecular event in Ewing sarcoma, leading to subsequent chromatin rearrangement and widespread transcriptional deregulation. Genomic profiling of chromatin accessibility shows that EWSR1::FLI1-bound regions become more accessible, with *GGAA* microsatellite binding frequently resulting in *de novo* enhancer formation (20). However, other studies have shown more limited, context-specific binding of EWSR1::FLI1 to chromatin (21, 22). In established Ewing sarcoma cells, binding at consensus ETS motifs frequently serves a repressive function by displacing wild-type ETS transcription factors from conserved enhancers (7). Notably, a zebrafish model of tumor initiation revealed a distinct mechanism, in which EWSR1::FLI1 reprogrammed neural crest cells toward a mesodermal fate and tumor initiation through binding at conserved ETS HA-containing developmental enhancers rather than GGAA microsatellites, which remained inaccessible in the earliest time points (23). Thus, it is important to understand the ability of EWSR1::FLI1 to recognize and bind nucleosome motifs and the molecular mechanisms by which it may do so.

Here, we investigated a key PF property - the ability of EWSR1::FLI1 to target different DNA motifs within nucleosomes - since PFs can occupy nucleosomal sites with an affinity similar to that of naked DNA. We prepared recombinant EWSR1::FLI1 and investigated its DNA and nucleosome affinities in biochemical and fluorescence-based experiments. To provide insight into the domains of EWSR1::FLI1 that contribute to its pioneering activities, we generated truncated FLI1 DBD constructs to assess their functionalities. We studied the targeting of these constructs to both the consensus HA motif and short *GGAA* microsatellites to determine the influence of binding motifs on the ability of EWSR1::FLI1 to target naked DNA and nucleosome motifs. Our results reveal that both the FLI1 DBD alone and full-length EWSR1::FLI1 can invade nucleosomes by trapping them in a partially unwrapped state at both the HA motif and short GGAA repeats. Interestingly, when compared to the FLI1 DBD alone, full-length EWSR1::FLI1 shows weakened binding affinity on naked DNA, but not on nucleosomal DNA motifs, with a shift to preferentially binding short *GGAA* repeats. Together, this work provides direct evidence that EWSR1::FLI1 possesses a key PF property in that it targets its DNA binding sites within nucleosomes similarly to naked DNA and that full-length EWSR1::FLI1 binding shifts to favor GGAA repeat targeting.

## Methods

### DNA Preparation

The Widom 601 sequence was cloned into the pUC19 plasmid and stored in the *Escherichia coli* DH5α strain. ETS consensus sequence (5’-*ACCGGAAGT*-3’) and GGAA microsatellites were inserted into the Widom 601 sequence via site-directed mutagenesis using the Site-Directed Mutagenesis Kit - QuikChange II (Agilent Technologies). DNA was amplified through polymerase chain reaction (PCR) using either Cy3-labeled oligonucleotides or non-labeled oligonucleotides (**Table S1**) that were obtained from Integrated DNA Technologies (IDT) and the *Pfu* DNA polymerase. Amplified DNA segments were purified through high-performance liquid chromatography (HPLC) using an anion-exchange column (either Mono Q 5/50 GL Ion Exchange Column, Cytiva Life Sciences or Gen-Pak FAX Anion-Exchange Chromatography Columns, Waters). Purified DNA segments were further buffer-exchanged and stored in 0.5x TE buffer (5 mM Tris–HCl, pH 8.0 and 0.5 mM ethylenediaminetetraacetic acid [EDTA]) using the Amicon Ultra Centrifugal Filter 30 kDa molecular weight cut off (MWCO) (Sigma-Aldrich). DNA constructs used for this study are shown in **Figure 1A**.

**Figure 1.**
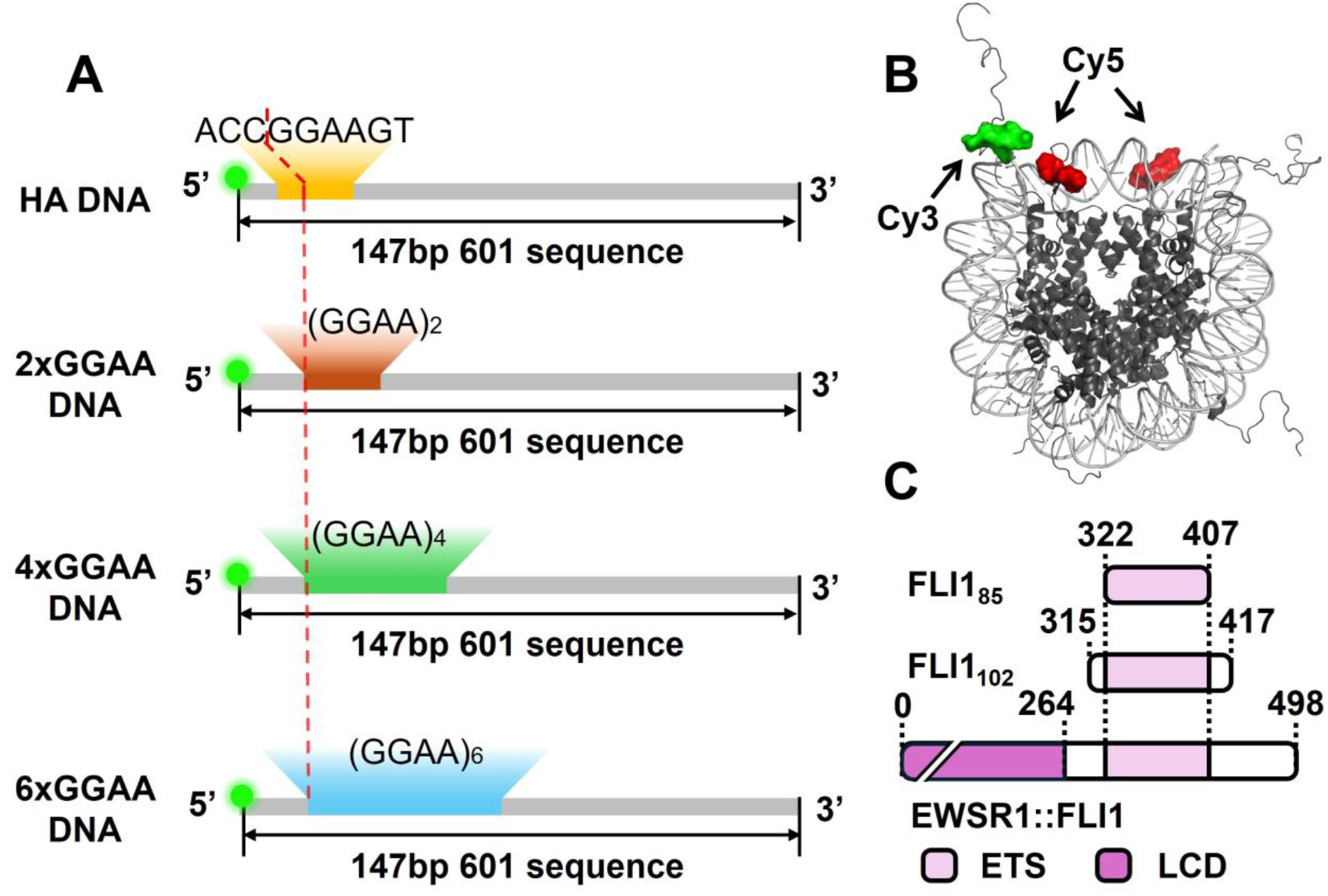
Schematic of DNA and protein constructs. **(A)** Illustration of DNA used for DNA binding experiments and nucleosome reconstitution. The Widom 601 sequence (grey) is used as NPS to yield homogenous recombinant nucleosomes, and the binding sites of EWSR1::FLI1 are inserted 8 or 10 bp away from the 5’ end of the DNA sequence. All DNA constructs are externally labeled at the 5’ end. **(B)** The nucleosome construct used in binding experiments (26). A Cy3 fluorophore is attached to the 5’ end of the nucleosomal DNA, and Cy5 fluorophores are labeled at H2AK119C within the histone octamer. **(C)** Diagrams of protein constructs used in binding experiments. FLI1_85_ includes the whole DBD (light purple), FLI1102 contains DBD with the flanking domains, and EWSR1::FLI1 is the full-length protein that includes the LCD domain (purple).

### Histone Octamer Preparation

The human histones (H2AK119C, H2B, and H4) and *Xenopus laevis* histones (H3C110A) used in these studies were obtained from The Histone Source (https://histonesource-colostate.nbsstore.net/). The H2A-H2B dimers and H3-H4 tetramers were refolded into the histone octamer in a 1.3:1 molar ratio through gradually dialyzing from the unfolding buffer (20 mM Tris-HCl pH 7.5, 7 M guanidine hydrochloride, 10mM dithiothreitol [DTT]) to the refolding buffer (10 mM Tris-HCl pH 7.5, 2 M NaCl, 1 mM EDTA pH 8.0, 5 mM β-mercaptoethanol [β-ME]) as previously reported (24). The refolded octamer was further labeled with Cy5 fluorophores at H2AK119C through the thiol-maleimide conjugation reaction. The labeling reaction was incubated at room temperature for an hour and continued for another 16 hours at 4 ⁰C before being quenched by 10 mM DTT. The labeled octamers were purified from excess H2A-H2B dimers through fast protein liquid chromatography (ÄKTA pure™ chromatography system, Cytiva) using a size-exchange chromatography (SEC) column (HiLoad S200 16/600 Superdex column, Cytiva). Purified octamers were concentrated using the Amicon Ultra Centrifugal Filter 30 kDa MWCO (Sigma-Aldrich) and added the same volume of 80% glycerol before storing in -20 ⁰C. The final product was examined through the sodium dodecyl sulfate–polyacrylamide gel electrophoresis (SDS-PAGE) with a 16% polyacrylamide gel and stained with Coomassie Brilliant Blue (**Figure S1A**).

### Nucleosome Preparation

DNA and histone octamers were reconstituted into recombinant nucleosomes in a 2:1 molar ratio through the salt dialysis method from a high salt condition (2 M NaCl, 5 mM Tris–HCl, pH 8.0, 0.5 mM EDTA, 1 mM benzylamine) to a low salt condition (5 mM Tris–HCl, pH 8.0, 0.5 mM EDTA, 1 mM benzylamine) as previously reported (25, 26). The recombinant nucleosomes were further purified through sucrose gradient (5-30%) ultracentrifugation method at 4 ⁰C, 41000 rpm, 22 hours on an ultracentrifuge (Beckman Coulter). Fractions after centrifugation were collected at a 0.4 mL volume and examined on a 5% acrylamide native gel. Fractions contained recombinant nucleosomes were further buffer-exchanged and stored in 0.5x TE buffer with 20% glycerol added before flash-freezing in liquid nitrogen and storage at -80 ⁰C. The final products were examined through running a 5% native acrylamide gel with 0.3x TBE buffer (30 mM Tris base, 27 mM boric acid, and 0.3 mM EDTA) at 300V for 1 hour under room temperature. The gels were imaged under Cy3 and Cy5 channel using a Typhoon biomolecular imager (**Figure S2A-B**).

### FLI185 and FLI1102 Preparation

FLI1_85_ (amino acids [a.a.] 322–407 of full-length Type I EWSR1::FLI1) and FLI1_102_ (a.a. 315–417 of full-length Type I EWSR1::FLI1) were expressed in *Escherichia coli* BL21(DE3) cells from pET28a expression vectors containing a C-terminal 6×His tag. Protein expression was induced with 1 mM IPTG at an OD600 of 1.0–1.2 and cultures were incubated overnight at 16 °C. Cells were harvested by centrifugation, resuspended in lysis buffer (300 mM KCl, 25 mM Bis-tris pH 6.9, 25 mM imidazole, 5 mM β-ME, and protease inhibitors (Roche 4693159001)), and lysed via sonication. The lysate was centrifuged at 10,000 × g for 30 min and the supernatant was incubated with Ni-NTA resin (Qiagen) for 1 h at 4 °C. Resin-bound protein was washed over a column with 90 mL of wash buffer (1M KCl, 25 mM Bis-tris, 40 mM imidazole, 5 mM β-ME, pH 6.9) then eluted with imidazole-containing buffer (600 mM KCl, 25 mM Bis-Tris, 400 mM imidazole, 5 mM β-ME, pH6.9). Eluted protein was dialyzed overnight into low-salt buffer (50 mM KCl, 25 mM Bis-Tris pH 6.5, 5 mM β-ME) and treated with nuclease (Pierce 88700).

The dialyzed proteins were further purified by cation-exchange chromatography using a 1mL HiTrap SP HP column (Cytiva). Proteins were eluted with a linear KCl gradient from 50 mM to 1 M KCl, and fractions containing purified FLI1_85_ or FLI1_102_ were identified by SDS-PAGE analysis (**Figure S1B-C**). Fractions containing the target protein were pooled, dialyzed into storage buffer (65 mM KCl, 25 mM Tris-HCl pH 7.9, 10% glycerol, 6 mM MgCl2, 0.5 mM EDTA, 0.2 mM Phenylmethlsulfonyl Fluoride [PMSF], and 1 mM DTT), concentrated using the Amicon Ultra Centrifugal Filter (Sigma Aldrich), flash-frozen in liquid nitrogen, and stored at −80 °C until use. A260/A280 ratios for purified proteins were determined to be between 0.55 and 0.58.

### EWSR1::FLI1 Preparation

Full-length EWSR1::FLI1 was expressed in *Escherichia coli* BL21(DE3) cells from a pET28a expression vector containing a C-terminal 6×His tag. Protein expression was induced with 1 mM IPTG at an OD600 of 0.5–0.6 and cultures were incubated for 3 h at 30 °C. Cells were harvested by centrifugation, resuspended in 20 mM Tris-HCl (pH 7.8) and 10 mM EDTA, supplemented with protease inhibitors (Roche 4693159001) and 1 mM PMSF, and lysed via sonication. The lysate was centrifuged at 10,000 × g for 20 min and the insoluble fraction was collected and washed 3 x 30 mL with wash buffer (20 mM Tris-HCl pH 7.8, 10 mM EDTA, and 1% Triton X-100). The washed pellet was solubilized in denaturing CAPS buffer (20 mM CAPS pH 11.0 and 6 M urea), clarified by centrifugation and filtration through a 0.45 µM syringe-driven filter (Cytiva Protein prep for AKTA systems). The solubilized fraction was incubated with Ni-NTA resin (Qiagen) for 2 h at 4 °C. Resin-bound protein was washed with CAPS buffer and eluted with CAPS buffer supplemented with 500 mM imidazole.

Eluted protein was refolded by dialysis against CAPS dialysis buffer (20 mM CAPS pH 11.0 and 10% glycerol) to remove urea. The refolded protein was concentrated using Amicon Ultra Centrifugal Filters (Sigma Aldrich) and further purified by SEC using a HiLoad Superdex 16/600 200 pg column (Cytiva) equilibrated in CAPS dialysis buffer (20 mM CAPS pH 11.0 and 10% glycerol). Fractions corresponding to EWSR1::FLI1 were identified by SDS-PAGE analysis (**Figure S1D**), pooled, concentrated as necessary, flash-frozen in liquid nitrogen, and stored at −80 °C until use.

### Electrophoresis Mobility Shift Assay (EMSA)

Both DNA- and nucleosome-binding events were monitored by EMSAs. We incubated 2 nM DNA or nucleosomes with a titrated amount of FLI1_85_, FLI1_102_, or full-length EWSR1::FLI1 in T130 buffer (130 mM NaCl, 10 mM Tris-HCl, pH 8.0, 10% glycerol, and 0.0075% Tween-20) for 20 minutes at room temperature. A 5% native acrylamide gel was prepared and ran in 0.3x TBE buffer. Gels were pre-run at 300 V for 90 minutes and were run at 300 V for another 90 minutes at 4 ⁰C. Gel shift results were obtained by imaging with a Typhoon biomolecular imager using the Cy3 or Cy5 channel as needed. Band intensity was further quantified and analyzed using the ImageJ (NIH) software. All measurements were complete with three replicates and binding isotherms were fit to the Hill equation to calculate S_1/2_. Binding specificity of FLI_85_, FLI_102_, and EWSR1::FLI1 toward naked DNA and nucleosomes were examined through performing EMSA measurements using 601 sequences without any binding motifs (**Figure S5A-F**).

### Förster Resonance Energy Transfer (FRET)

To perform FRET assays, we labeled 5’ end of our DNA constructs with Cy3 fluorophore (donor) as described above (**Figure 1A-B**) and the nucleosome with the Cy5 fluorophore (acceptor) at H2AK119C (**Figure 1B**). Titrations of FLI1_85_, FLI1_102_, or full-length EWSR1::FLI1 were incubated with 2 nM nucleosomes for 20 minutes at room temperature in T130 buffer. After incubation, fluorescence spectra were immediately obtained by FluoroMax 4 fluorometer (Horiba). We excited Cy3 fluorophore at 510 nm and Cy5 fluorophore at 610 nm and respectively measured their emission from 530 to 750 nm and 630 to 670 nm. FRET efficiency was calculated using the ratio_A_ method, as previously reported (27). All of the measurements were completed with three replicates. The result of the binding between nucleosomes and protein constructs was fit to the Hill equation to calculate S_1/2_.

## Results

### The FLI1 DBD targets consensus ETS motifs in nucleosomes with similar efficiency as fully exposed DNA

Large-scale genomics approaches indicate that oncogenic gene regulation in Ewing sarcoma requires EWSR1::FLI1 to function as a PF (7, 20, 21, 28). EWSR1::FLI1 binds both high-affinity ETS sites and GGAA microsatellites in cells, and putative PF activity at both classes of motifs are implicated in establishing aberrant gene regulation in Ewing sarcoma (29–31). Since PFs are the first TFs that target a silenced gene, a key property of PFs is their ability to target DNA sites wrapped within nucleosomes similarly to fully exposed DNA sites (8, 32). Previous studies show that PFs target their sites within nucleosomes at affinities that are within a factor of 10 relative to fully exposed DNA sites, while the occupancy of canonical TFs can be suppressed by 100- to 10,000-fold (10, 13, 33–35). In addition, based on prior studies of other PFs showing that the intrinsically disordered regions of TFs have a limited effect on pioneer activity (10, 33), we started with a minimalist approach to establish the baseline DNA and nucleosome binding properties of the structured FLI1 DBD. While the FLI1 DBD is typically defined as an 85-a.a. wHTH (FLI1_85_, **Figure 1C**), our recent work identified a larger 102-a.a. DBD that includes an additional C-terminal α4 helix (FLI1_102_, **Figure 1C**) as the minimal functionally oncogenic DBD for EWSR1::FLI1 (36, 37). We therefore investigated the nucleosome and DNA binding properties of both FLI1_85_ and FLI1_102_ to understand the role of the FLI1 wHTH in nucleosome targeting and of the α4 helix in this binding activity.

Other ETS TFs, like PU.1, have demonstrated PF activity on ETS motifs in the nucleosome entry-exit region (38). We therefore focus on this region of the nucleosome. We used the 147 bp Widom 601 nucleosome positioning sequence (NPS) with a 9-bp ETS HA motif (5’-*ACCGGAAGT*-3’) inserted at 8 - 16 bp from the 5’ end of the left side of the 601 NPS (**Figure 1A**). This positions the HA motif near the DNA entry-exit site of the nucleosome. For binding studies with DNA only, we separately prepared Widom 601 DNA with the same HA motif positioning within the 601 NPS as that in our nucleosomes (**Figure 1A**). Quantification of the DNA and nucleosome constructs was done by detecting the fluorescence from the Cy3 fluorophore attached to the 5’ end of the DNA or the Cy5 fluorophore attached to the nucleosome at H2AK119C, respectively (**Figure 1B**). Binding events were monitored by EMSAs and analyzed by determining the fraction of unshifted DNA or nucleosome band to calculate the value of S_1/2_. The S_1/2_ is the concentration at which half of the DNA or nucleosomes are bound and is approximately the apparent dissociation constant (apparent K_D_). We observed both FLI1_85_ and FLI1_102_ binding to the HA site-containing DNA, as evidenced by slower-migrating protein-DNA complexes within EMSA gels (**Figure 2A** and **2B**). Quantification reveals that FLI1_85_ (S_1/2_ = 2.3 ± 0.2 nM) and FLI1_102_ (S_1/2_ = 2.7 ± 0.2 nM) bind to HA site-containing DNA with similar nanomolar binding affinity (**Figure 2E and Table S2**).

**Figure 2.**
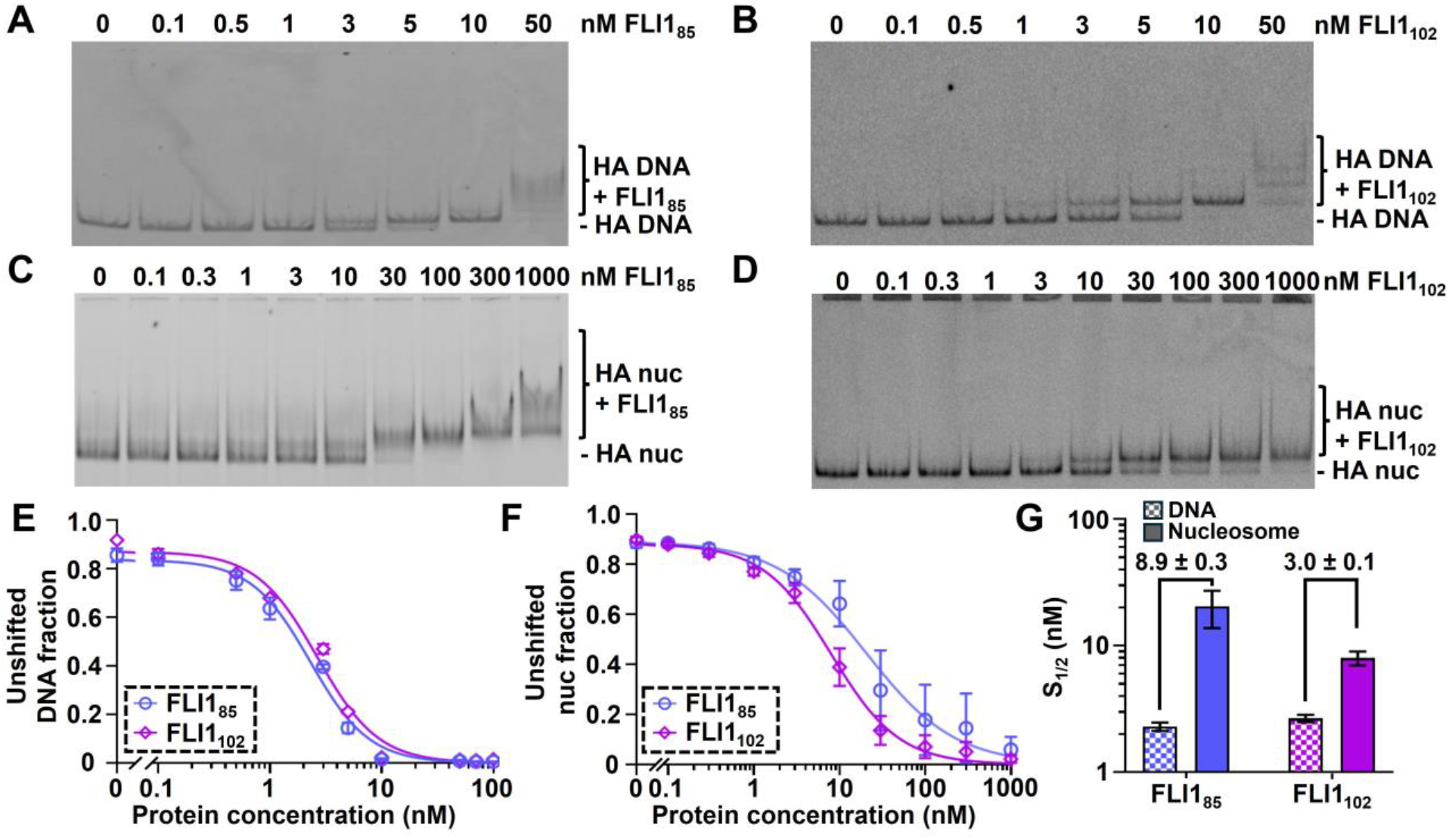
EWSR1::FLI1 DBD binds DNA and nucleosomes with an HA sequence. **(A)** EMSA of FLI185 binding to free 601 DNA with HA site. The DNA band is completely shifted when binding to 10 nM FLI185. The gel is imaged with the Cy3 channel. **(B)** EMSA of FLI1102 binding to free DNA with HA motif. The shifting of the DNA band starts to occur at 1 nM FLI1102, and the band is completely shifted when binding to 10 nM FLI1102. The gel is scanned with the Cy3 channel. **(C)** EMSA binding of FLI185 to nucleosomes with an HA site. The nucleosome band is completely shifted upward when binding to 100 nM FLI185. The gel is imaged with the Cy5 channel. **(D)** EMSA of FLI1102 binding to nucleosomes containing an HA motif around the entry-exit region. The shifting first occurs at 3 nM FLI1102 and completes at 1000 nM FLI1102. The EMSA gel is scanned with the Cy5 channel. **(E)** Quantification results of EMSA measurement of FLI185 and FLI1102 binding to DNA containing an HA site. **(F)** Quantification results of FLI185 and FLI1102 binding to nucleosomes contained HA motif near the entry-exit region. **(G)** Summary of S_1/2_ for the FLI185 and FLI1102 binding measurements in panel E and F. FLI185 binding to DNA with HA site: 2.3 ± 0.2 nM. FLI1102 binding to DNA with HA site: 2.7 ± 0.2 nM. FLI185 binding to nucleosomes with HA site: 20 ± 7 nM. FLI1102 binding to nucleosome with HA site: 8 ± 1 nM. The fold change in S_1/2_ when comparing nucleosome to DNA binding is shown above each set of bars. All EMSA measurements were performed in triplicates.

We then characterized the affinity of FLI1_85_ and FLI1_102_ to nucleosomes containing the HA site. EMSAs show that both FLI1_85_ and FLI1_102_ interact with HA site-containing nucleosomes to form a protein-nucleosome complex (**Figure 2C** and **Figure 2D**). Quantification of the EMSAs reveals that FLI1_85_ (S_1/2_ = 20 ± 7 nM) and FLI1_102_ (S_1/2_ = 8 ± 1 nM) bind nucleosomes within a factor of 10 of their affinities to DNA (**Figure 2F** and **Figure 2G**; 8.9 ± 0.3-fold for FLI1_85_ and 3.0 ± 0.1-fold for FLI1_102_). These results indicate that both FLI1_85_ and FLI1_102_ exhibit the pioneering property that nucleosomes do not efficiently block their occupancy at HA motifs. Notably, FLI1_102_ exhibits a smaller difference in binding affinities between nucleosomes and DNA compared to FLI185 (3.0 ± 0.1-fold vs. 8.9 ± 0.3-fold), suggesting that the flanking C-terminal α4 helix of the FLI1 DBD enhances nucleosome binding (**Figure 2G**). Together, these results indicate that the wHTH of the EWSR1::FLI1 DBD can efficiently target the HA ETS motif in both DNA and nucleosomes, with the flanking residues present in FLI1_102_ enhancing nucleosome-binding.

### The FLI1 DBD targets Ewing sarcoma response elements in nucleosomes with similar efficiency as fully exposed DNA

In addition to the ETS consensus sequence, repetitive GGAA microsatellites are critical EWSR1::FLI1 response elements in Ewing sarcoma, with 4-6 GGAA repeats comprising the minimal repeat length needed for transcriptional activation (15, 29). Recent structural evidence further supports that the truncated FLI1 DBD can cooperatively bind to GGAA repeats (39). building off our finding that FLI185 and FLI1102 show efficient nucleosome binding at consensus ETS motifs, we tested whether this activity is also seen with short repetitive GGAA sequences. As before, we used EMSAs to monitor DNA and nucleosome binding. Here we used three short GGAA repeats – 2xGGAA, 4xGGAA, and 6xGGAA (**Figure 1A**) – to investigate the length dependence within GGAA repeats that comprise the minimal motif required for EWSR1::FLI1-mediated transcriptional activation in *in vitro* studies (15).

We first focused on FLI185 and FLI1102 binding to 147 bp DNA that contained the 601 NPS with GGAA repeats inserted, where the first GGAA was at the same position as the HA core GGAA (**Figure 1A**). Our EMSA gels show that both FLI185 and FLI1_102_ bind to DNA containing 2xGGAA repeats and form DNA-protein complexes with slower gel mobilities, leading to multiple shifted bands with FLI1 DBD concentrations above 10 nM (**Figure 3A and 3B).** Similar electromobility shifts are observed for FLI1_85_ and FLI1_102_ binding to DNA containing either 4xGGAA or 6xGGAA repeats (**Figure S3A-B** and **Figure S4A-B**). Quantification of these EMSAs indicates that the length of GGAA repeats does not significantly influence the binding affinity of FLI1_85_ and FLI1_102_ (**Figure 3E**, **S3E**, **S4E**, and **Table S2**; S_1/2 2xGGAA DNA-FLI1_85 = 4.3 ± 0.2 nM, S_1/2 4xGGAA DNA-FLI1_85 = 5.6 ± 0.9 nM, S_1/2 6xGGAA DNA-FLI1_85 = 5.5 ± 0.8 nM, S_1/2 2xGGAA DNA-FLI1_102 = 5.9 ± 0.8 nM, S_1/2 4xGGAA DNA-FLI1_102 = 6.7 ± 0.9 nM, S_1/2 6xGGAA DNA-FLI1_102 = 5.8 ± 0.5 nM). The occurrence of multiple shifted bands when titrating with FLI1_85_ and FLI1_102_ suggests multiple binding events by the FLI1 DBD to GGAA microsatellites, consistent with the prior published evidence using the truncated FLI1 wHTH (39).

**Figure 3.**
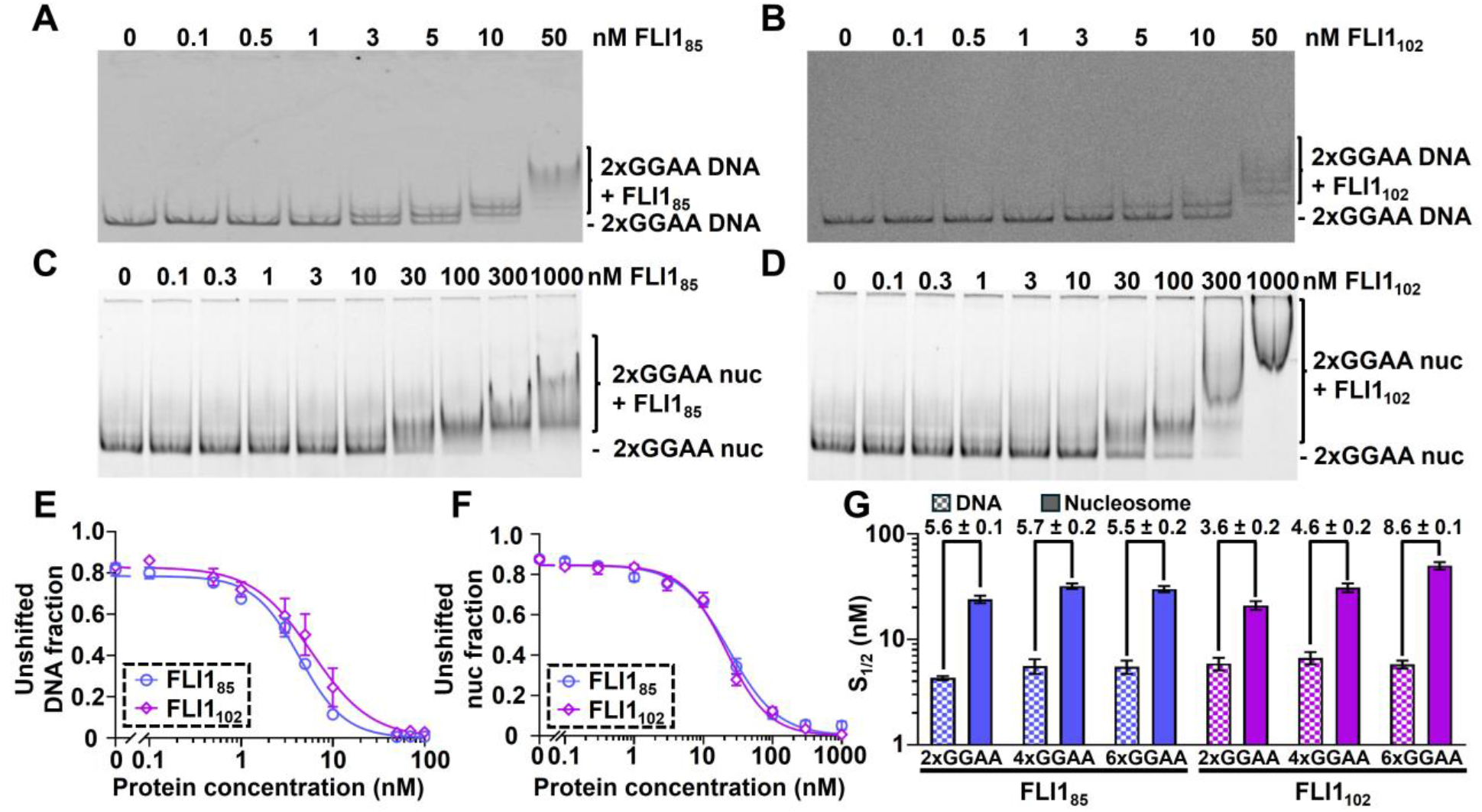
EWSR1::FLI1 DBD binds DNA and nucleosomes with short GGAA repeats. **(A)** EMSA of FLI185 binding to Cy3-labeled free 601 DNA inserted with 2xGGAA microsatellites. The image is scanned with the Cy3 channel. **(B)** EMSA of FLI1102 binding to free 601 DNA containing 2xGGAA repeats. The image is scanned with the Cy3 channel. **(C)** EMSA of FLI185 binding to nucleosomes containing 2xGGAA microsatellites. The image is scanned with the Cy5 channel. **(D)** EMSA of FLI1102 binding to nucleosomes containing 2xGGAA microsatellites. The image is scanned with the Cy5 channel. **(E)** Quantification results of EMSA measurements of FLI185 and FLI1102 binding to free 601 DNA with a 2xGGAA motif. **(F)** Quantification results of EMSA measurements of FLI1_85_ and FLI1102 binding to nucleosomes containing 2xGGAA sequences near the entry-exit region. **(G)** Summary of S_1/2_ values for FLI185 and FLI1102 binding to DNA and nucleosomes containing 2xGGAA, 4xGGAA, and 6xGGAA microsatellites. FLI185 binding to DNA with 2xGGAA repeat: 4.3 ± 0.2 nM. FLI1102 binding to DNA with 2xGGAA repeat: 5.9 ± 0.8 nM. FLI185 binding to DNA with 4xGGAA repeat: 5.6 ± 0.9 nM. FLI1102 binding to DNA with 4xGGAA repeat: 6.7 ± 0.9 nM. FLI1_85_ binding to DNA with 6xGGAA repeat: 5.5 ± 0.8 nM. FLI1102 binding to DNA with 6xGGAA repeat: 5.8 ± 0.5 nM. FLI185 binding to nucleosomes with 2xGGAA repeat: 24 ± 2 nM. FLI1102 binding to nucleosomes with 2xGGAA repeat: 21 ± 2 nM. FLI1_85_ binding to nucleosomes with 4xGGAA repeat: 32 ± 2 nM. FLI1102 binding to nucleosomes with 4xGGAA repeat: 31 ± 3 nM. FLI185 binding to nucleosomes with 6xGGAA repeat: 30 ± 2 nM. FLI1102 binding to nucleosomes with 6xGGAA repeat: 50 ± 3 nM. The fold change in S_1/2_ when comparing nucleosome to DNA binding is shown above each set of bars. All EMSA measurements were performed in triplicates.

We then investigated the efficiency of FLI1_85_ and FLI1_102_ binding to nucleosomes containing GGAA motifs by repeating our nucleosome binding EMSA experiments with nucleosomes containing the 2xGGAA, 4xGGAA, and 6xGGAA DNA constructs. We found that both FLI1_85_ and FLI1_102_ bind GGAA repeat-containing nucleosomes around 20 to 50 nM (**Figure 3C-D, S3C-D,** and **S4C-D**). EMSA quantification shows that FLI1_85_ (S_1/2 2xGGAA Nuc-FLI1_85 = 24 ± 2 nM) and FLI1_102_ (S_1/2 2xGGAA Nuc-FLI1_102 = 21 ± 2 nM) bind 2xGGAA-containing nucleosomes with similar affinities (**Figure 3F**). Interestingly, FLI1_85_ has similar binding affinities for 4xGGAA and 6xGGAA (S_1/2 4xGGAA Nuc-FLI1_85 = 32 ± 2 nM, S_1/2 6xGGAA Nuc-FLI1_85 = 30 ± 2 nM), while FLI1_102_ nucleosome binding weakens as the GGAA repeat length increases (S_1/2 4xGGAA Nuc-FLI1_102 = 31 ± 3 nM, S_1/2 6xGGAA Nuc-FLI1_102 = 50 ± 3 nM) (**Figure 3F**, **, S3F, S4F, and Table S2**).

To determine if FLI1_85_ and FLI1_102_ target GGAA repeat-containing nucleosomes efficiently as it does for HA site-containing nucleosomes, we compared S_1/2_ values between DNA and nucleosome that contain the same GGAA repeat. For FLI1_85_, the S_1/2_ values for binding nucleosomes relative to DNA were consistently ∼5-fold higher at all three GGAA lengths (5.6 ± 0.1-fold at 2xGGAA, 5.7 ± 0.2-fold at 4xGGAA, and 5.5 ± 0.2-fold at 6xGGAA). In contrast, for FLI1102, the S_1/2_ values for nucleosome binding relative to DNA increased as the GGAA repeat number increased (3.6 ± 0.2-fold, 4.6 ± 0.2-fold, and 8.6 ± 0.1-fold for 2xGGAA, 4xGGAA, and 6xGGAA, respectively, **Figure 3G**). Importantly, these results show that both FLI185 and FLI1102 bind to nucleosomes relative to fully exposed DNA with less than a 10-fold difference in affinity, implying they retain the efficient nucleosome targeting pioneering property at GGAA repeats. In addition, we find that including the C-terminal α4 helix with the FLI1 DBD results in less efficient nucleosome binding as the GGAA repeat number increases up to six repeats.

### The FLI1 DBD can invade and trap nucleosomes in partially unwrapped states

(40) Previous studies of PFs that bind near the entry-exit site of the nucleosome, including the ETS factor PU.1, show that PF binding to this region of the nucleosome traps them in a partially unwrapped state (10, 38, 41). Having shown that the FLI1 DBD, with and without the C-terminal α4 helix, can efficiently bind within the nucleosome entry-exit region at both HA and GGAA repeat motifs with EMSAs, we next investigated whether FLI185 and FLI1102 occupancy at these motifs trapped nucleosomes in partially unwrapped states. To do this, we used an ensemble FRET assay to monitor nucleosome partial unwrapping (10, 42–44). Nucleosomes were prepared with a donor fluorophore (Cy3) attached to the 5’ end of the DNA adjacent to the binding motif, and an acceptor fluorophore (Cy5) attached at H2AK119C (**Figure 1B**). These fluorophore positions result in high FRET efficiency for fully wrapped nucleosomes, whereas nucleosomes in partially unwrapped states result in significantly lower FRET efficiency.

We first separately titrated FLI185 and FLI1102 with nucleosomes that contained the HA motif and measured the fluorescence emission spectra when exciting Cy3 at 515 nm **(Figure 4A upper panel**) and Cy5 at 610 nm **(Figure 4A lower panel**). When exciting Cy3 with 515 nm without either FLI1 DBD, there is a significant Cy5 fluorescence peak at about 650 nm, which decreases as either FLI185 or FLI1102 is included at increasing concentrations (**Figure 4A, S7A**). In addition, as the Cy5 emission peak decreases, the Cy3 emission increases as expected for a decrease in FRET efficiency (**Figure 4A, S7A**). We determined the FRET efficiency for each FLI185 and FLI1102 concentration using the ratio_A_ method (27) (**Figure 4B**) and found that the reduction in FRET efficiency changes over similar ranges of concentration. The FRET efficiency versus FLI185 and FLI1102 concentration fit to binding isotherms with S_1/2_ values (S_1/2 HA Nuc-FLI1_85 = 47 ± 6 nM and S_1/2 HA Nuc-FLI1_102 = 38 ± 6 nM; **Table S2**) that are within 2-fold and 5-fold of the S_1/2_ for nucleosome binding in the EMSA measurements, respectively (**Table S3**). These 2- and 5-fold differences are similar to those observed with other TFs (10, 13) and are likely due to EMSA measurements being inherently non-equilibrium, while the FRET measurements are in equilibrium. We confirmed that without the HA binding motif, the FRET efficiency did not decrease with increasing FLI1 DBD concentrations, which demonstrates that the reduction in FRET is due to FLI1 DBD binding to the HA motif (**Figure S7**).

**Figure 4.**
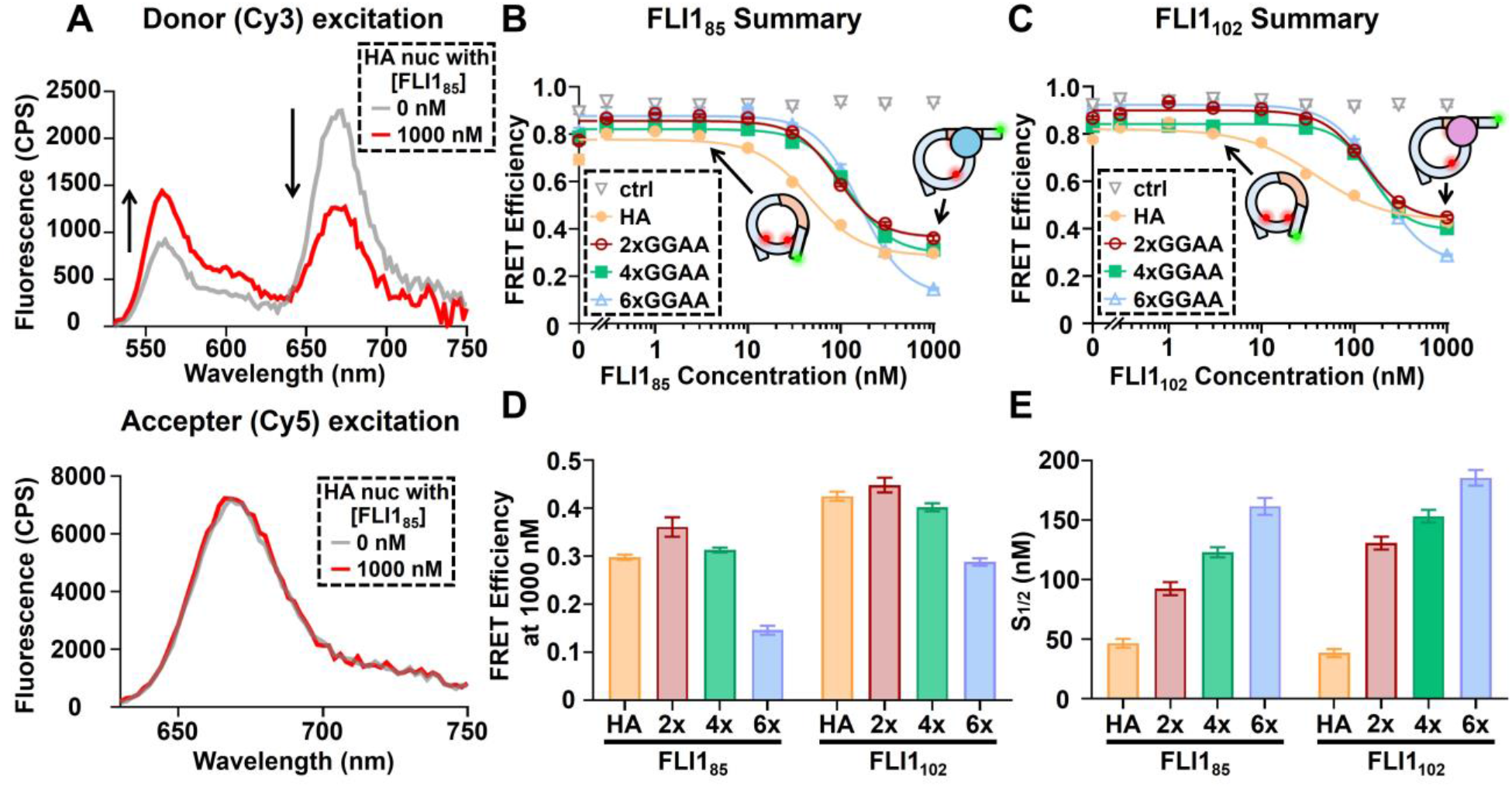
The binding of EWSR1::FLI1 DBD to nucleosomes induces partial nucleosome unwrapping. **(A)** Representative fluorescence spectrum of 2 nM nucleosomes containing a HA site binding to 0 nM FLI185 (grey) and 1000 nM FLI185 (red). **(B)** ΔFRET of nucleosomes binding to a titrated amount of FLI185. **(C)** ΔFRET of nucleosomes binding to a titrated amount of FLI1102. **(D)** The value of FRET efficiency when nucleosomes containing various GGAA motifs bind to 1000 nM FLI185 or FLI1_102_. **(E)** The value of S_1/2_ obtained from panel B and C. FLI185 binds to nucleosomes with HA motif at an S_1/2_ of 47 ± 6 nM, 2xGGAA repeats at an S_1/2_ of 92 ± 9 nM, 4xGGAA repeats at an S_1/2_ of 123 ± 7 nM, and 6xGGAA repeats at an S_1/2_ of 160 ± 10 nM. FLI1102 binding to nucleosomes with HA motif at an S_1/2_ of 38 ± 6 nM, 2xGGAA microsatellite at an S_1/2_ of 131 ± 9 nM, 4xGGAA microsatellite at an S_1/2_ of 153 ± 9 nM, and 6xGGAA microsatellite at an S_1/2_ of 190 ± 10 nM. All FRET measurements were done in triplicate and the error bar is generated from the standard error.

We then investigated if FLI1_85_ and FLI1_102_ trap nucleosomes in partially unwrapped states when targeting GGAA repeat motifs. As we did for the EMSA measurements, we carried out FRET efficiency measurements with nucleosomes containing 2x, 4x, and 6xGGAA repeats, where the first GGAA is in the same location as the HA motif. As with the HA motif, we find that the reduction in FRET fits to binding isotherms for both FLI1_85_ and FLI1102 binding to all three GGAA repeat lengths (**Figure 4B-C, S6A-C, S7B-D**). The S_1/2_ values (S_1/2 2xGGAA Nuc-FLI1_85 = 92 ± 9 nM, S_1/2 4xGGAA Nuc-FLI1_85 = 123 ± 7 nM, S_1/2 6xGGAA Nuc-FLI1_85 = 160 ± 10 nM, S_1/2 2xGGAA Nuc-FLI1_102 = 131 ± 9 nM, S_1/2 4xGGAANuc-FLI1_102 = 153 ± 9 nM, S_1/2 6xGGAA Nuc-FLI1_102 = 190 ± 10 nM; **Table S2**) are comparable but about 3 to 6 times higher than the EMSA determined S_1/2_ of nucleosome binding (**Figure 4E**, **Table S3**). Comparison of these results to FLI185 and FLI1102 binding to nucleosomes without a GGAA motif implies that the reduction in FRET involves site specific binding of FLI185 and FLI1102 to trap partially unwrapped nucleosomes. Thus, as seen for the HA motif, the differences in S_1/2_ values between the EMSA and FRET measurements for short GGAA repeats could arise from differences in the assays with EMSAs measuring a non-equilibrium state and FRET measuring the system at equilibrium. Interestingly, the 6x GGAA motif resulted in a larger reduction in FRET efficiency, which is consistent with additional FLI1 DBDs binding to the longer motif, which extends further into the nucleosome (**Figure 4D**). Overall, these results reveal that both FLI1_85_ and FLI1102 can trap nucleosomes in a partially unwrapped state, which could help open chromatin to facilitate transcription activation.

### The relative affinity of EWSR1::FLI1 between DNA motifs reverses, with preferential binding to GGAA repeats over the consensus motif

Having established the nucleosome binding properties of the minimal truncated FLI1 DBD constructs, we next sought to understand the nucleosome binding properties of full-length, wild-type EWSR1::FLI1. Full-length EWSR1::FLI1 contains additional disordered regions from the C-terminus of FLI1 flanking the DBD as well as the intrinsically disordered LCD from the N-terminus of EWSR1 (45). Previous studies of EWSR1::FLI1 binding to DNA have either focused on just the FLI1 portion of the fusion protein for quantitative analysis (17, 29) or have been limited to semi-quantitative biochemical or cell-based genomics assays (7, 15, 16, 20, 21, 29, 36, 46, 47). While these studies show there is a difference in DNA and chromatin binding specificity for the fusion protein, the quantitative and mechanistic basis for this difference remains unclear. To better understand how the DNA and nucleosome binding properties of full-length, wild-type EWSR1::FLI1 compare to those of the minimal DBD, we developed an approach for recombinant expression of full-length, wild-type EWSR1::FLI1 (See Methods for details). Our approach resulted in soluble biochemical quantities of EWSR1::FLI1 in native buffer conditions.

After obtaining full-length EWSR1::FLI1, we investigated its binding affinity to DNA at the HA site and short GGAA repeats with EMSAs. We found that EWSR1::FLI1 bound to all motifs with a significantly lower affinity compared to the minimal DNA domain. Interestingly, full-length EWSR1::FLI1 binds DNA containing the consensus HA site with a significantly lower affinity as compared to short GGAA repeats (**Figure 5A**). We observed that DNA containing each of the GGAA repeats is nearly fully bound by EWSR1::FLI1 at 300 nM, while DNA containing the HA motif is largely unbound with 300 nM EWSR1::FLI1 (**Figure 5A**). EMSA quantification shows that the EWSR1::FLI1 S_1/2_ with GGAA repeats (S_1/2 2xGGAA Nuc-EWSR1::FLI1_ = 96 ± 5 nM, S_1/2 4xGGAA Nuc-EWSR1::FLI1_ = 94 ± 5 nM, S_1/2 6xGGAA Nuc-EWSR1::FLI1_ = 100 ± 10 nM; **Figure 5B-C**, **Table S2**) is about an order of magnitude smaller than with the HA motif (S_1/2 HA Nuc-EWSR1::FLI1_ > 600 nM; **Figure 5B-C**, **Table S2**). These results indicate that full-length EWSR1::FLI1 prefers short GGAA repeats over the consensus HA ETS site by about an order of magnitude. Importantly, given our previous finding that EWSR1::FLI1 has a binding stoichiometry of ∼1 EWSR1::FLI1 molecule for every 2 GGAA repeats (17) these results indicate that altered sequence specificity emerges at motifs that bind only to a single EWSR1::FLI1 (i.e. the HA and 2xGGAA motifs). Notably, the DNA-binding preference of EWSR1::FLI1 is reversed from either FLI185 or FLI1102, suggesting that other domains of EWSR1::FLI1 may alter sequence-specific binding by the FLI1 DBD.

**Figure 5.**
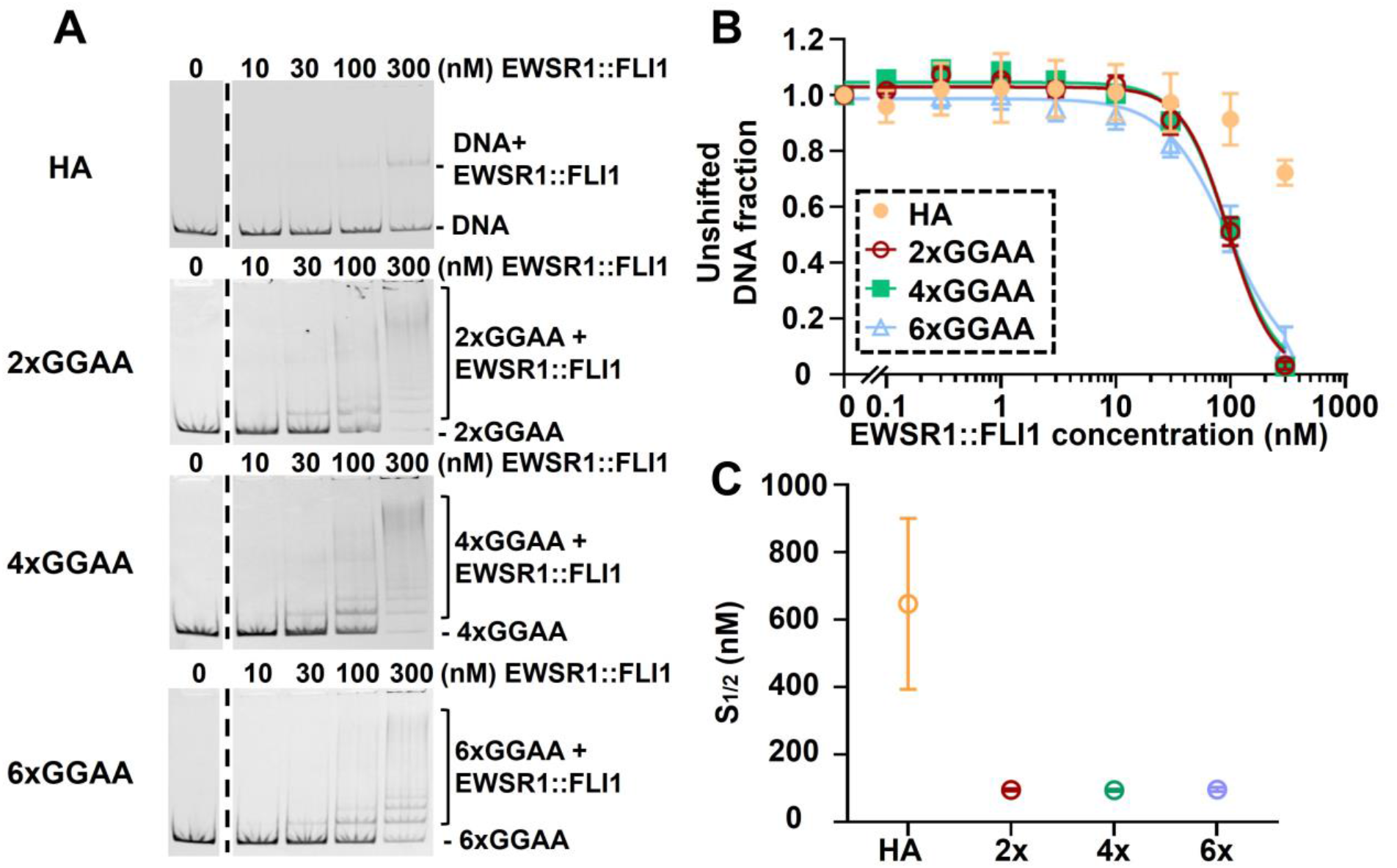
EWSR1::FLI1 binds to HA-inserted DNA and GGAA-inserted DNA with distinct affinities. **(A)** Representative EMSA gels of EWSR1::FLI1 binding to free 601 DNA containing different motifs - HA site, 2xGGAA repeats, 4xGGAA repeats, and 6xGGAA repeats. The EMSA images are scanned with the Cy3 channel. **(B)** Quantification results of EMSA measurements in panel A. **(C)** Summary of S_1/2_ values obtained from fitting the binding isotherm of panel **C** to the Hill equation. EWSR1::FLI1 binding to free 601 DNA with a HA site at an S_1/2_ of 600 ± 400 nM, 2xGGAA microsatellite at an S_1/2_ of 96 ± 5 nM, 4xGGAA microsatellite at an S_1/2_ of 94 ± 5 nM, and 6xGGAA microsatellite at an S_1/2_ of 100 ± 10 nM. All EMSA measurements were completed in triplicate. The error bar indicates the standard error.

### EWSR1::FLI1 can invade and trap nucleosomes in partially unwrapped states

Given our observation that EWSR1::FLI1 preferentially recognizes GGAA repeats within fully exposed DNA, we investigated EWSR1::FLI1 binding to nucleosomes containing these binding motifs. We carried out EMSAs of nucleosomes containing either GGAA repeats or the HA motif with increasing concentrations of EWSR1::FLI1 (**Figure 6A**). We find that EWSR1::FLI1 binds nucleosomes with a significantly larger S_1/2_ than both FLI185 and FLI1102, which is consistent with the significantly larger S_1/2_ to DNA. We find that EWSR1::FLI1 binds HA-motif-containing nucleosomes at around 500 nM and GGAA-repeat-containing nucleosomes at around 600-1000 nM. This implies that EWSR1::FLI1 binds the HA motif within nucleosomes similarly to fully exposed DNA, while it binds GGAA repeats within nucleosomes with about a 10-fold weaker affinity than DNA. These results indicate that while EWSR1::FLI1 binds both nucleosomes and DNA with lower affinities to the FLI DBD domain alone, it retains the pioneering property of binding its target sites within nucleosomes with similar affinities to fully exposed DNA.

**Figure 6.**
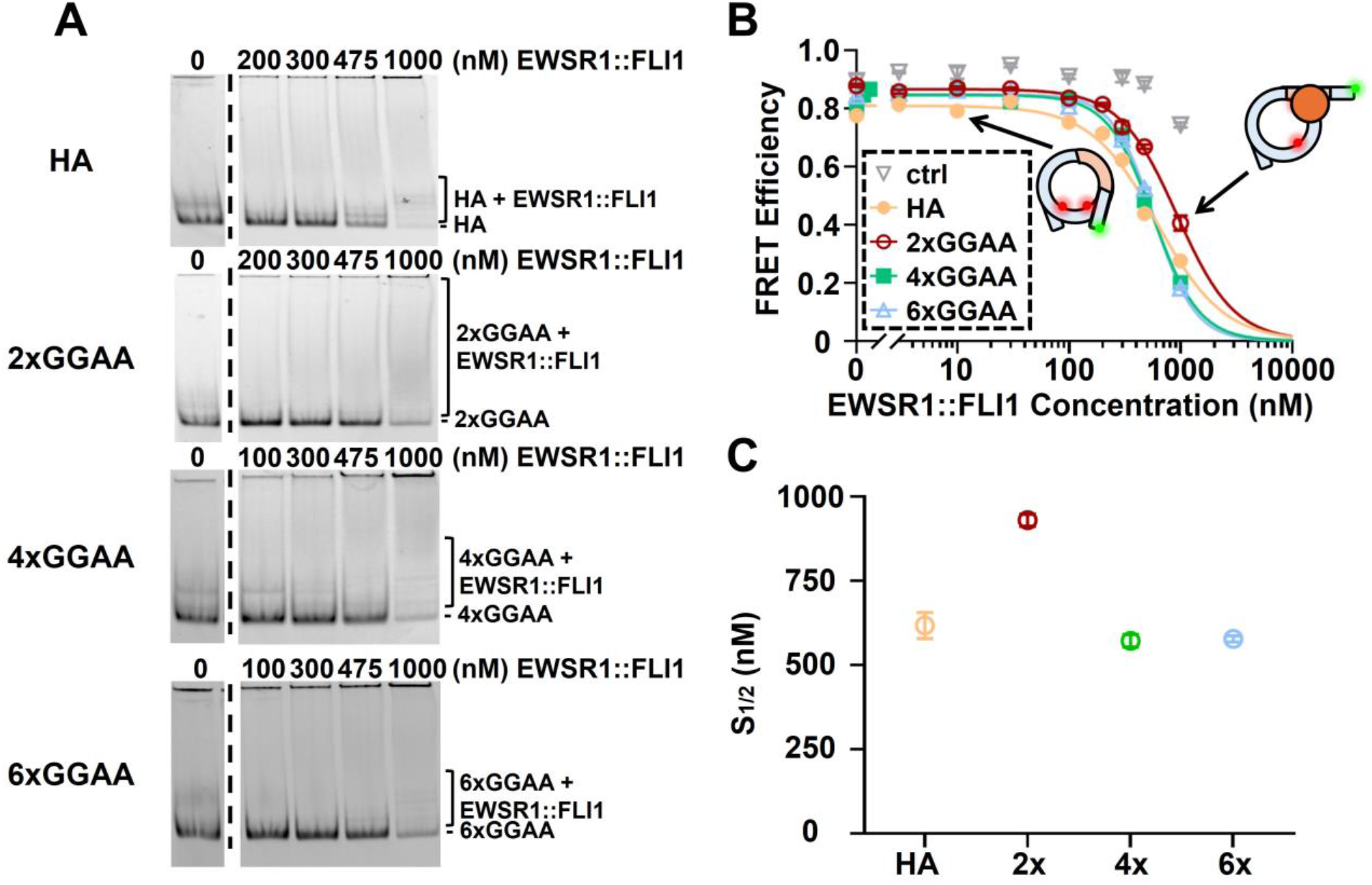
EWSR1::FLI1 binds to nucleosome-embedded HA site and GGAA microsatellites, altering nucleosome structure. **(A)** Representative EMSA gels of EWSR1::FLI1 binding to nucleosomes containing various motifs around the entry-exit region, including HA site, 2xGGAA, 4xGGAA, and 6xGGAA microsatellites. The EMSA gels are scanned with the Cy5 channel. All EMSA gels were performed in triplicate. **(B)** Ensemble FRET measurements of EWSR1::FLI1 binding to nucleosomes. A fixed concentration of nucleosome (2nM) was incubated with a titrated amount of EWSR1::FLI1. The illustrated cartoon indicates how EWSR1::FLI1 binding decreases FRET efficiency. The Hill equation was used to fit the binding isotherms. All measurements were completed in triplicate. **(C)** Summary of S_1/2_ obtained from the FRET measurements. EWSR1::FLI1 binding to nucleosomes with an HA site at an S_1/2_ of 620 ± 40 nM, 2xGGAA microsatellite at an S_1/2_ of 930 ± 20 nM, 4xGGAA microsatellite at an S_1/2_ of 570 ± 20 nM, and 6xGGAA microsatellite at an S_1/2_ of 580 ± 10 nM.

Given that EWSR1::FLI1 also exhibits the pioneering property of efficient nucleosome binding, we investigated if this binding traps nucleosomes in partially unwrapped states, as we observed for FLI1 DBD alone. To do this, we relied on FRET measurements of the Cy3-Cy5 labeled nucleosomes (**Figure 1B**) with either the HA motif or the GGAA repeats near the nucleosome entry-exit region, as we did with the FLI1 DBD constructs. We find that titrations of EWSR1::FLI1 result in a reduction in the FRET efficiency over the concentration ranges that EMSAs detect nucleosome binding (**Figure 6B, S8A-D**). The FRET efficiency measurements of EWSR1::FLI1 titrations fit to binding isotherms with S_1/2_ values between 500 -1000 nM (**Figure 6C**; S_1/2 HA Nuc-EWSR1::FLI1_ = 620 ± 40 nM, S_1/2 2xGGAA Nuc-EWSR1::FLI1_ = 930 ± 20 nM, S_1/2 4xGGAA Nuc-EWSR1::FLI1_ = 570 ± 20 nM, S_1/2 6xGGAA Nuc-EWSR1::FLI1_ = 580 ± 10 nM; **Table S2**). Importantly, without a binding motif, the reduction in FRET efficiency was much smaller, demonstrating that the impact on nucleosome unwrapping is largely motif-dependent (**Figure 6B**). Overall, these results show that full-length EWSR1::FLI1 can efficiently bind within nucleosomes when targeting either the HA motif or GGAA repeats.

## Discussion

In this study, we leverage our ability to prepare not only truncated EWSR1::FLI1 DBD constructs, but also full-length EWSR1::FLI1, to determine that they target both the HA motif and GGAA repeats within the entry-exit region of nucleosomes with a less than 10-fold change in efficiency relative to naked DNA. This effect was the most pronounced for EWSR1::FLI1 targeting its HA site where it appears to bind nucleosomes better than DNA. In addition, FLI1102 was extremely efficient at targeting nucleosomes at HA sites with a 3-fold lower binding affinity relative to DNA alone (**Figure 2G, 5C, 6C**). These data suggest that the C-terminal α4 helix promotes nucleosome engagement by the DBD at consensus ETS motifs. Consistent with previous binding studies of human PFs, our results reveal that EWSR1::FLI1, and FLI1102 are efficient at targeting their sites within nucleosomes (51). For comparison, motif occupancy of non-pioneering TFs such as Pho4 and Gal4 at the same location in the entry-exit region of the nucleosome as studied here are inhibited by nucleosome structure by 100 to 10,000-fold (13, 33), whereas established PFs generally bind DNA and nucleosomes with similar affinities (10, 13). For example, PU.1, a well-known ETS-family PF, has only 8-fold difference in affinities between free and nucleosome DNA (51). This pattern is consistent with the dissociation rate compensation mechanism, which is a mechanism for how PFs efficiently target their binding motifs within nucleosomes. For this PF mechanism, a reduced dissociation rate from nucleosomes relative to DNA alone compensates for a reduced binding rate constant to its site within nucleosomes (10). Together, these studies provide significant quantitative evidence that both EWSR1::FLI1 and its minimal DBD possess pioneering activity.

Another PF property is that PF binding within chromatin increases accessibility for other transcription activators to bind their sites. Here, our FRET measurements indicate that the binding of both truncated and full-length EWSR1::FLI1 traps nucleosomes into partially unwrapped states through the site exposure model (52), which can increase the probability of other TFs binding events (43). Notably, the longer GGAA 6x repeat resulted in greater nucleosome unwrapping (**Figure 4D, 6B**), suggesting that for even longer repeats more DNA could be trapped in an unwrapped state. In addition to nucleosome unwrapping, PFs also can facilitate nucleosome repositioning alone, when in the context of native NPS. For example, PU.1 shifts the *CX3CR1* nucleosome by 17 bp (37), ASCL1/E12α shift the *NRCAM* nucleosome in a stepwise process (54), and the budding yeast PF Reb1 can shift nucleosomes on native NPSs (53). These observations suggest that EWSR1::FLI1 binding at longer GGAA repeats may not only unwrap, but also reposition and even destabilize nucleosomes, which has been observed for other PFs and TF binding on repetitive motifs (55–57). Future studies will be required to investigate these additional mechanisms.

This study relied on the strong Widom 601 NPS because it enabled preparation of homogeneously positioned nucleosomes. This allowed us to investigate nucleosome substrates where the position of the motifs is well-defined and PF targeting nucleosomes occurs via the site exposure model. Also, since the partially unwrapping of Widom 601 NPS is similar to native NPSs, we focused on motifs positioned near the nucleosome entry/exit site (54). Both motif position and native NPSs will influence PF targeting and its impact on nucleosome position and stability. Motifs closer to the dyad will have different accessibility, while longer GGAA repeats near the dyad may decrease nucleosome stability. Having established the ability to prepare full-length EWSR1::FLI1, future studies using nucleosomes containing native NPS with different motif positions will be important. In addition, since chromatin is the *in vivo* target of EWSR1::FLI1, future studies within nucleosome arrays will also be critical for understanding the impact of EWSR1::FLI1 on chromatin compaction.

GGAA repeats are some of the most studied EWSR1::FLI1 response elements, whereas EWSR1::FLI1 function at consensus ETS motifs is less understood and likely context-specific. While HA sites have recently been highlighted as playing an essential role in tumor initiation (23), EWSR1::FLI1 binding at HA sites has also been implicated in EWSR1::FLI1-mediated repression (7). According to our biochemical studies, both FLI185 and FLI1102 exhibit a lower S_1/2_ at the HA site compared to any of the GGAA repeats when binding to its motif in free DNA and to partially unwrapped nucleosomes (**Figure 2** and **Figure 4E**), which is consistent with strong, sequence-specific binding to the full 9 bp HA motif. In contrast, full-length EWSR1::FLI1 binds GGAA repeats more efficiently than the HA site, even on the 2xGGAA repeat, which has a similar binding stoichiometry as the HA site (17) (one EWSR1::FLI1 molecule bound per site, **Figure 5**). This shift in sequence preference from the HA site to even short GGAA repeats in free DNA has not been previously observed in biochemical studies of the FLI1 DBD (29). Notably, nucleosomes suppressed full-length EWSR1::FLI1 binding to the 2xGGAA site by ∼10-fold, while binding to the HA site was not suppressed, suggesting that histones may interact with EWSR1::FLI1 in a sequence-specific manner to promote HA site binding (**Figure 6**). Together, these differences in binding preference indicate that distinct sequence-specific binding modes at different DNA motifs may shape the oncogenic activity of EWSR1::FLI1 in chromatin. Additionally, future studies are needed to further determine how histone tails or histone modifications further modulate EWSR1::FLI1 recruitment, as is suggested by previous cell-based studies (22).

Although DBD is the domain that directly interacts with the targeted binding motif, other domains within the TF, especially the intrinsically disordered regions (IDR), play important roles in regulating the binding events. Both *in vivo* and *in vitro* studies have suggested that EWSR1 LCD triggers phase separation and can form biomolecular condensates by promoting homo- and heterotypic intermolecular self-association interactions (45, 58–60). Previous studies have also reported that EWSR1 LCD facilitates transcriptional hub formation at GGAA microsatellites in the cellular context (59, 61, 62). Formation of these types of assemblies could change the kinetics of DNA and nucleosome binding. Conversely, binding to DNA has been shown to impair condensate formation by EWSR1::FLI1 (60). Our results show that the EWSR1 LCD and intrinsically disordered regions (IDR) of FLI1 confer EWSR1::FLI1 distinct sequence-specific binding in free DNA compared to the truncated wHTH FLI1 DBD. Future experiments will be important to determine if the engagement of nucleosomes affects EWSR1::FLI1 condensate formation and whether formation of condensates modifies the pioneering activity of EWSR1::FLI1.

Finally, genome regulation in Ewing sarcoma is characterized by EWSR1::FLI1 binding at GGAA microsatellites, formation of *de novo* enhancers, and downstream transcriptional activation (7, 63). Recent work using genetically engineered models further suggests that EWSR1::FLI1 engagement of consensus ETS motifs is important for tumor initiation by EWSR1::FLI1 in neural crest cells (23). Oncogenic transcriptional activation at these loci relies on the ability of EWSR1::FLI1 to bind and function at its target motifs in chromatin, requiring pioneering activity to gain access to DNA in nucleosomes. By determining a recombinant purification approach to prepare full-length, soluble EWSR1::FLI1, we quantitatively defined how EWSR1::FLI1 and its minimal DBD bind to DNA and interact with nucleosomes at both HA sites and GGAA repeats for the first time. This new approach is well-positioned to enable a range of future mechanistic studies of EWSR1::FLI1 oncogenic functions.

## Acknowledgements

We thank Dr. Stephen Lessnick and Michelle Cruz for their support and enthusiasm for this project, as well as the Theisen and Poirier labs for valuable discussion regarding this project.

## Author Contributions

Ruo-Wen Chen (Conceptualization [equal], Formal analysis [lead], Investigation [lead], Validation [lead], Writing – original draft [lead], Writing – review and editing [equal]), Runwei Zhou (Formal analysis [supporting], Investigation [supporting], Writing – original draft [supporting], Writing - review and editing [supporting]), E. John Tokarsky (Conceptualization [equal], Formal analysis [supporting], Investigation [supporting]), Megann A. Boone (Conceptualization [equal], Formal analysis [supporting], Investigation [supporting]), Andrea K. Byrum (Conceptualization [equal], Formal analysis [supporting], Investigation [supporting]), Michael G. Poirier (Conceptualization [equal], Formal Analysis [supporting], Funding acquisition [supporting], Investigation [supporting], Project administration [supporting], Supervision [equal], Writing – original draft [supporting], Writing – review and editing [equal]), Emily R. Theisen (Conceptualization [equal], Formal Analysis [supporting], Funding acquisition [lead], Investigation [supporting], Project administration [lead], Supervision [equal], Writing – original draft [supporting], Writing – review and editing [lead]).

## Conflict of Interest

E.R.T. has previously received funding from Salarius Pharmaceuticals unrelated to the submitted work. The authors declare no additional conflicts of interest.

## Declaration of generative AI and AI-assisted technologies in the writing process

Generative AI and AI-assisted technologies were not used in the drafting or editing of this manuscript.

## Funding

E.R.T. was funded by a St. Baldrick’s Scholar Award, and Alex’s Lemonade Stand R-Accelerated Award, and the National Institutes of Health (R37 CA299623). M.G.P was supported with funding from the National Institutes of Health (R35 GM139654). R.Z. was supported by a CancerFree KIDS New Idea Award. The Typhoon imager for taking fluorescence images in this study were obtained through the support from the National Institutes of Health S10OD023582. Funding to pay the open access publication charges were provided by the National Institutes of Health R37 CA299623.

## Data Availability

Lead contact: Correspondence and requests should be addressed to Emily Theisen.

All data needed to evaluate and reproduce the results in the paper are present in the paper and/or the Supplmentary Materials. Sequences for synthetic oligonucleotides used in EMSA and FRET assays are detailed in relevant sections of the Methods, Figure 1, and Supplementary Table S1. Unique and stable reagents are available from Emily Theisen upon request and a completed materials transfer agreement with Nationwide Children’s Hospital.

