## Supplementary Figures and Tables for "Fusion Oncoprotein EWSR1::FLI1 Invades Nucleosomes at Consensus ETS Motifs and GGAA Microsatellites"

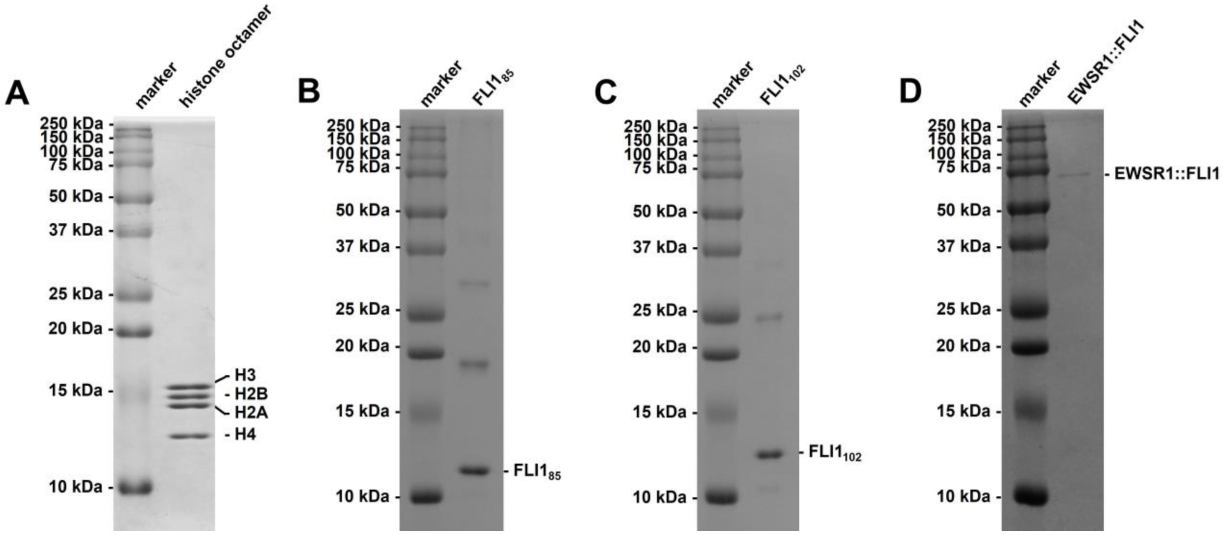


**Figure S1. Protein preparation for histone octamer, truncated EWSR1::FLI1 DBDs (FLI1$_{85}$ and FLI1$_{102}$), and full-length EWSR1::FLI1.** SDS-PAGE was performed to examine the result of protein preparation, including histone octamer **(A)**, truncated EWSR1::FLI1 (FLI1$_{85}$ **(B)**; FLI1$_{102}$ **(C)**), and full-length EWSR1::FLI1. The left lane was loaded with protein ladder, while the right lane was loaded with the target protein. The gels were stained with Coomassie brilliant blue.

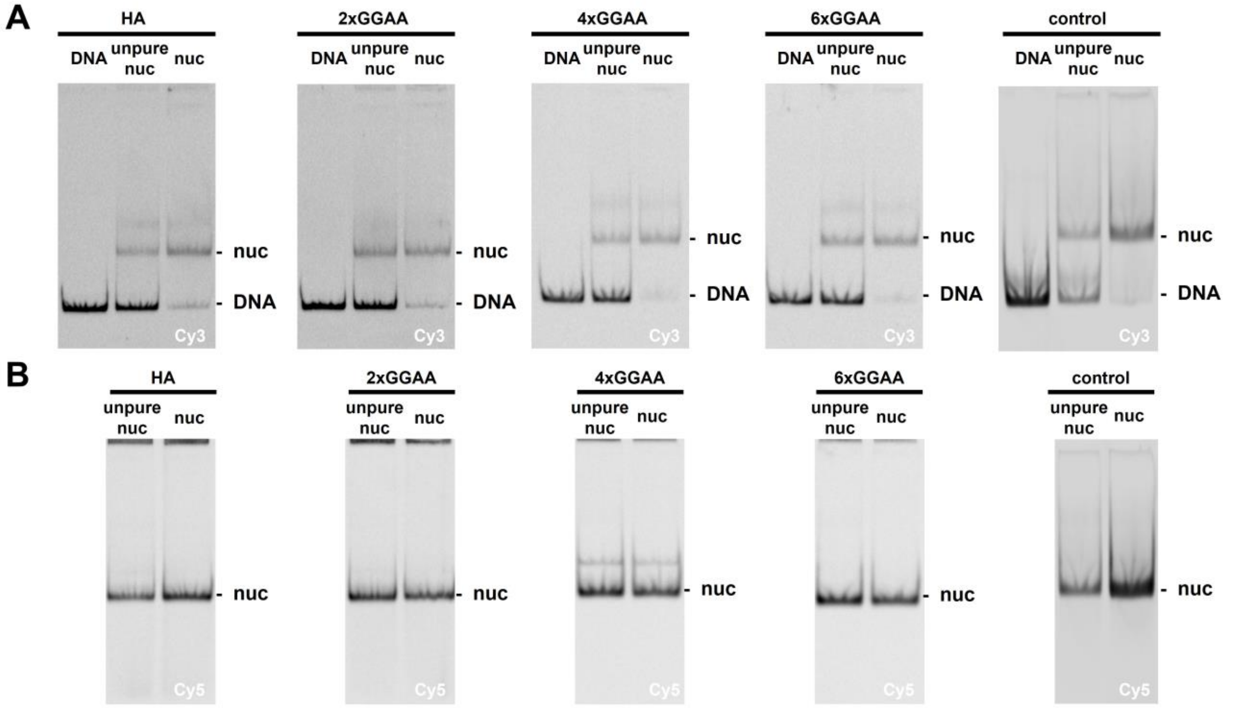


**Figure S2. Preparation of recombinant nucleosomes.** Excess DNA was removed by sucrose gradient centrifugation to obtain purified nucleosomes. The result of nucleosome reconstitution was examined by native gel electrophoresis with a 5% acrylamide gel. Each gel was loaded with purified DNA (the left lane), nucleosomes before purification (the middle lane, abbreviated as ”unpure nuc”), and nucleosome after purification (the right lane, abbreviated as ”nuc”). The gel was imaged by a Typhoon imager with the Cy3 channel for Cy3-labeled DNA **(A)** and the Cy5 channel for the Cy5-labeled histone octamer **(B)**.

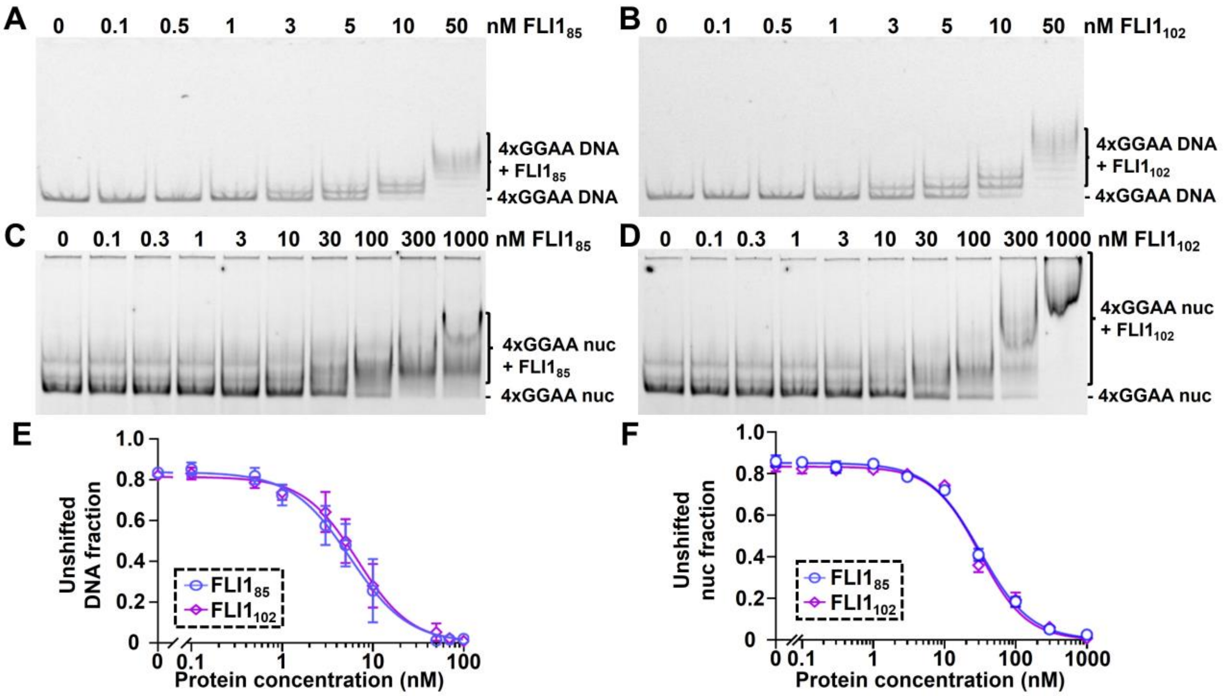


**Figure S3. Truncated EWSR1::FLI1 DBDs bind to 4xGGAA-inserted DNA/nucleosomes. (A,B)** EMSA of $FLI1_{85}$ **(A)** and $FLI1_{102}$ **(B)** binding to free 601 DNA with 4xGGAA repeats. The 5% native acrylamide gel is imaged with the Cy3 channel to probe Cy3-labeled DNA. The 4xGGAA DNA band starts to shift when binding to 3 nM $FLI1_{85}$ and $FLI1_{102}$. Multiple bands start to appear when the concentration of $FLI1_{85}$ and $FLI1_{102}$ is above 5 nM. **(C,D)** EMSA of $FLI1_{85}$ **(C)** and $FLI1_{102}$ **(D)** binding to nucleosome-embedded 4xGGAA motif. The 4xGGAA sequence begins at 10th base pair from the 5' end of DNA, which is around the entry-exit site of nucleosomes. The recombinant nucleosome is Cy5-labeled at H2AK119C. The 5% acrylamide native gel is scanned with the Cy5 channel by a Typhoon imager. The nucleosome band starts to shift upward when binding to 30 nM $FLI1_{85}$ or $FLI1_{102}$ and completely shifts when the concentration of $FLI1_{85}$ or $FLI1_{102}$ reaches 300 nM. **(E,F)** Quantification results of $FLI1_{85}$ and $FLI1_{102}$ binding to free 601 DNA **(E)** and nucleosome **(F)** with 4xGGAA repeats. All measurements were done in triplicate, and the error bar is generated from the standard error.

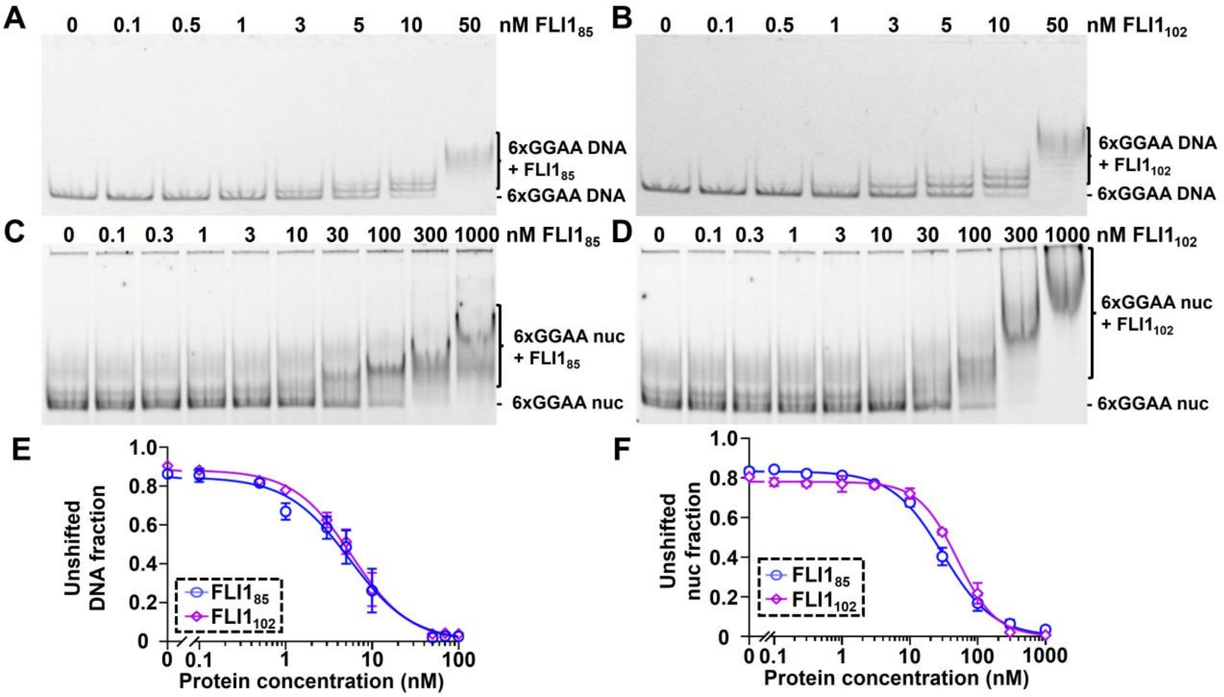


**Figure S4. Truncated EWSR1::FLI1 DBDs bind to 6xGGAA-inserted DNA/nucleosomes. (A,B)** EMSA of FLI1$_{85}$ **(A)** and FLI1$_{102}$ **(B)** binding to free 601 DNA with 6xGGAA repeats. The 5% native acrylamide gel is imaged with the Cy3 channel to probe Cy3-labeled DNA. The 6xGGAA DNA band starts to shift when binding to 3 nM FLI1$_{85}$ and FLI1$_{102}$. Multiple bands start to appear when the concentration of FLI1$_{85}$ and FLI1$_{102}$ is above 5 nM. **(C,D)** EMSA of FLI1$_{85}$ **(C)** and FLI1$_{102}$ **(D)** binding to nucleosome-embedded 6xGGAA motif. The 6xGGAA sequence begins at 10th base pair from the 5' end of DNA, which is around the entry-exit site of nucleosomes. The recombinant nucleosome is Cy5-labeled at H2AK119C. The 5% acrylamide native gel is scanned with the Cy5 channel by a Typhoon imager. The nucleosome band starts to shift upward when binding to 30 nM FLI1$_{85}$ or FLI1$_{102}$ and completely shifts when the concentration of FLI1$_{85}$ or FLI1$_{102}$ reaches 300 nM. **(E,F)** Quantification results of FLI1$_{85}$ and FLI1$_{102}$ binding to free 601 DNA **(E)** and nucleosome **(F)** with 6xGGAA repeats. All measurements were done in triplicate, and the error bar is generated from the standard error.

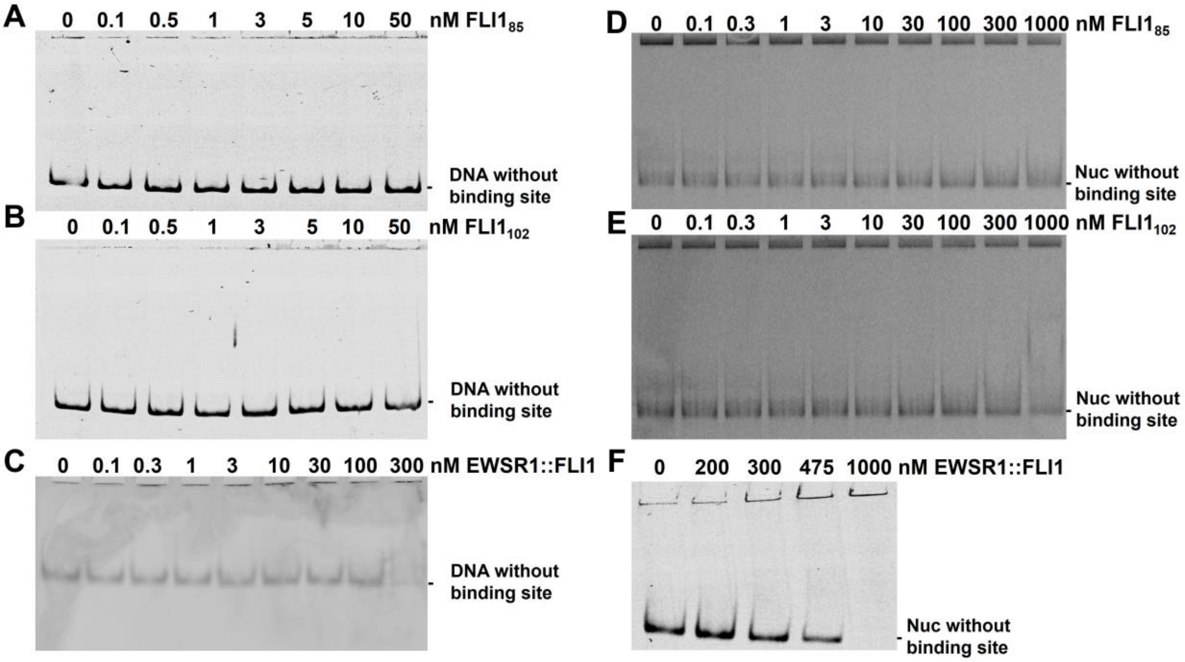


**Figure S5. Recombinant FLI1$_{85}$, FLI1$_{102}$, and EWSR1::FLI1 bind either DNA or nucleosome with specificity.** The binding experiments with 601 DNA or nucleosomes lacking any GGAA binding-motifs are shown here to examine non-specific binding. **(A)** EMSA of FLI1$_{85}$ binding to 601 147 bp DNA without any binding motifs. The measurement was imaged through the Cy3 channel to probe Cy3-labeled DNA. The reaction concentration of DNA is 2 nM. **(B)** EMSA of FLI1$_{102}$ binding to 601 147 bp DNA without any binding motifs. The measurement was imaged through the Cy3 channel to probe Cy3-labeled DNA. The reaction concentration of DNA is 2 nM. **(C)** EMSA of EWSR1::FLI1 binding to 601 147 bp DNA without any binding motifs. The measurement was imaged through the Cy3 channel to probe Cy3-labeled DNA. The reaction concentration of DNA is 2 nM. **(D)** EMSA of FLI1$_{85}$ binding to 601 147 bp nucleosomes without any binding motifs. The measurement was imaged through the Cy5 channel to probe Cy5-labeled histone octamers. The reaction concentration of nucleosomes is 2 nM. **(E)** EMSA of FLI1$_{102}$ binding to 601 147 bp nucleosomes without any binding motifs. The measurement was imaged through the Cy5 channel to probe Cy5-labeled histone octamers. The reaction concentration of nucleosomes is 2 nM. **(F)** EMSA of EWSR1::FLI1 binding to 601 147 bp nucleosomes without any binding motifs. The measurement was imaged through the Cy5 channel to probe Cy5-labeled histone octamers. The reaction concentration of nucleosomes is 2 nM.

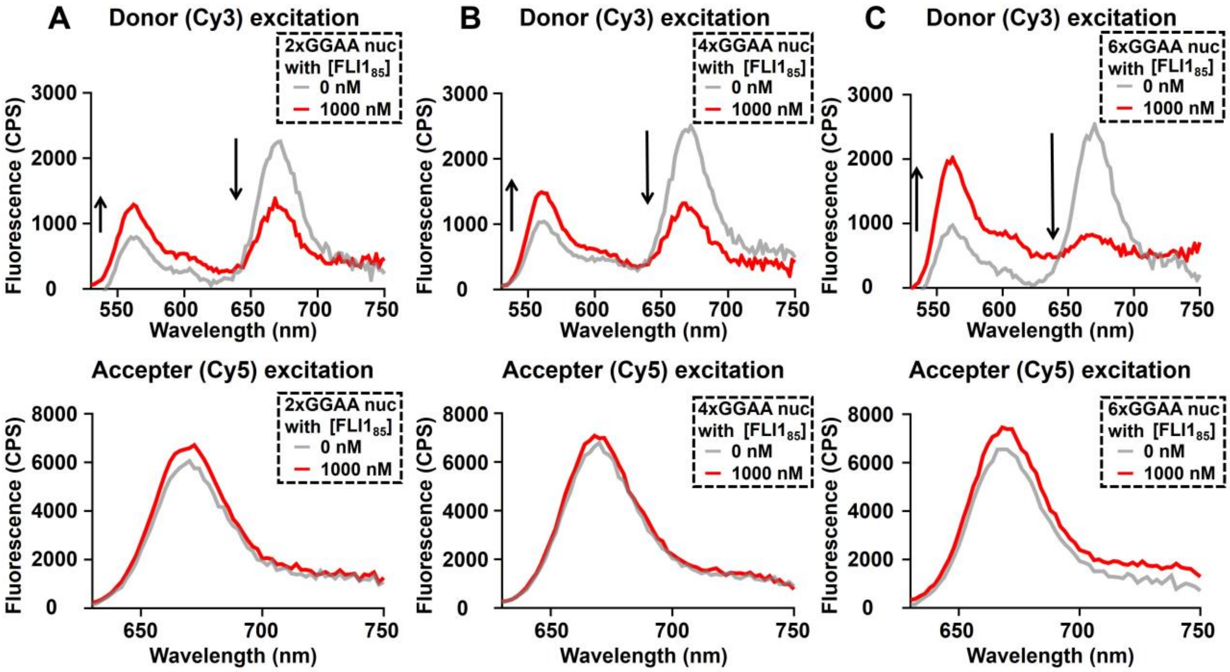


**Figure S6. FLI1$_{85}$ induces FRET changes in nucleosomes encoded with different lengths of GGAA repeats. (A)** FRET efficiency decreases after 2xGGAA-encoded nucleosomes were incubated with 1000 nM FLI1$_{85}$. The fluorescence spectrum is obtained from a fluorometer. Cy3 is excited at the upper panel, while Cy5 is excited at the lower panel. **(B)** FRET efficiency decreases after 4xGGAA-encoded nucleosomes were incubated with 1000 nM FLI1$_{85}$. The fluorescence spectrum is obtained from a fluorometer. Cy3 is excited at the upper panel, while Cy5 is excited at the lower panel. **(C)** FRET efficiency decreases after 6xGGAA-encoded nucleosomes were incubated with 1000 nM FLI1$_{85}$. The fluorescence spectrum is obtained from a fluorometer. Cy3 is excited at the upper panel, while Cy5 is excited at the lower panel.

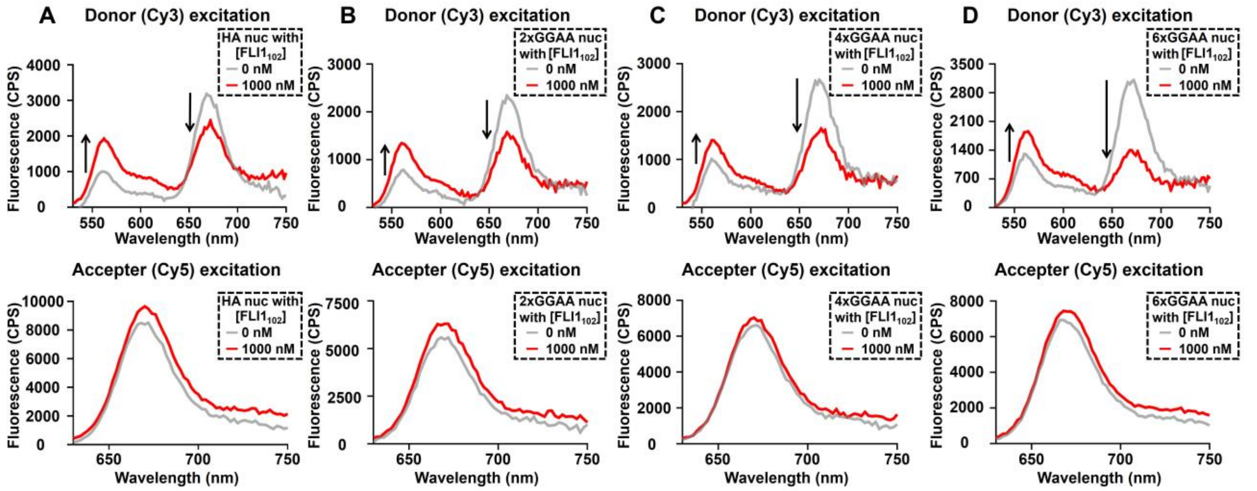


**Figure S7. FLI$_{102}$ induces FRET changes in nucleosomes encoded with ETS consensus sequence (HA) and different lengths of GGAA repeats. (A)** FRET efficiency decreases after HA nucleosomes were incubated with 1000 nM FLI1$_{102}$. The fluorescence spectrum is obtained from a fluorometer. Cy3 is excited at the upper panel, while Cy5 is excited at the lower panel. **(B)** FRET efficiency decreases after 2xGGAA-inserted nucleosomes were incubated with 1000 nM FLI1$_{102}$. The fluorescence spectrum is obtained from a fluorometer. Cy3 is excited at the upper panel, while Cy5 is excited at the lower panel. **(C)** FRET efficiency decreases after 4xGGAA-inserted nucleosomes were incubated with 1000 nM FLI1$_{102}$. The fluorescence spectrum is obtained from a fluorometer. Cy3 is excited at the upper panel, while Cy5 is excited at the lower panel. **(D)** FRET efficiency decreases after 6xGGAA-inserted nucleosomes were incubated with 1000 nM FLI1$_{102}$. The fluorescence spectrum is obtained from a fluorometer. Cy3 is excited at the upper panel, while Cy5 is excited at the lower panel.

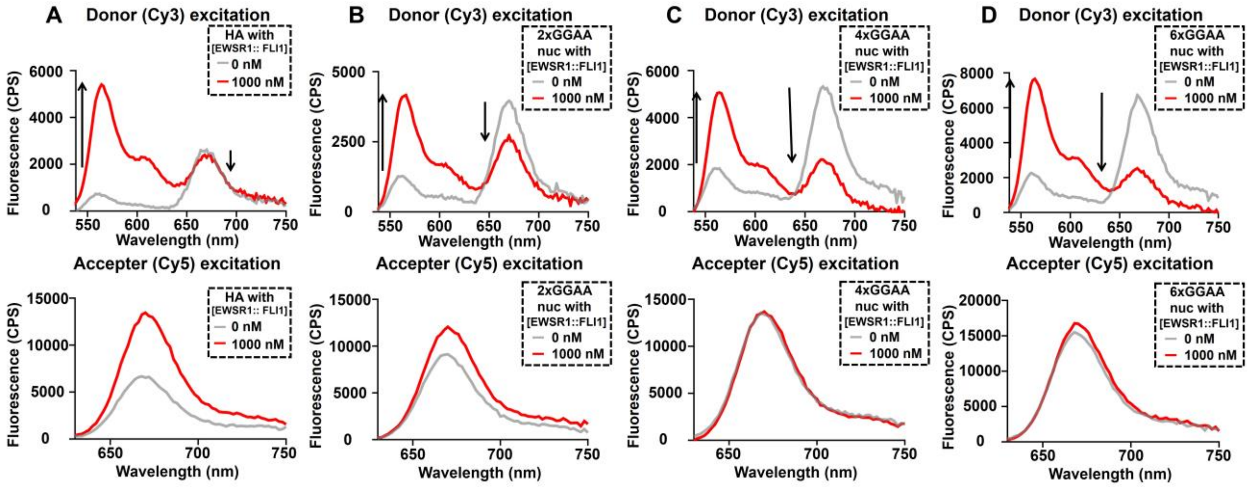


**Figure S8. Full-length EWSR1::FLI1 induces FRET changes in nucleosomes encoded with ETS consensus sequence (HA) and different lengths of GGAA repeats. (A)** FRET efficiency decreases after HA nucleosomes were incubated with 1000 nM EWSR1::FLI1. The fluorescence spectrum is obtained from a fluorometer. Cy3 is excited at the upper panel, while Cy5 is excited at the lower panel **(B)** FRET efficiency decreases after 2xGGAA-inserted nucleosomes were incubated with 1000 nM EWSR1::FLI1. The fluorescence spectrum is obtained from a fluorometer. Cy3 is excited at the upper panel, while Cy5 is excited at the lower panel. **(C)** FRET efficiency decreases after 4xGGAA-inserted nucleosomes were incubated with 1000 nM EWSR1::FLI1. The fluorescence spectrum is obtained from a fluorometer. Cy3 is excited at the upper panel, while Cy5 is excited at the lower panel**. (D)** FRET efficiency decreases after 6xGGAA-inserted nucleosomes were incubated with 1000 nM EWSR1::FLI1. The fluorescence spectrum is obtained from a fluorometer. Cy3 is excited at the upper panel, while Cy5 is excited at the lower panel.

**Table S1: Oligonucleotides used in the study**

| | |
|---|---|
| Primers for binding experiments (HA site in bold: **ACCGGAAGT**) | |
| Cy3-601 147bp For. | Cy3- CTGGAGAATCCCGGTGCC |
| 601 147bp Rev. | ACAGGATGTATATATCTGACACG |
| Cy3-HA 147bp For. | Cy3- CTGGAGA**ACCGGAAGT**CC |
| Cy3-GGAA 147bp For. | Cy3- CTGGAGAATCGGAAGG |
| Primers for cloning through site-directed mutagenesis | |
| EWSHighAffinityFwd | CCAATTGAGCGGCCTCGGACTTCCGGTTCTCCAG GAATTCACTGGCCGTCGTTTTACA |
| EWSHighAffinityRevs | TGTAAAACGACGGCCAGTGAATTCCTGGAGAACCG GAAGTCCGAGGCCGCTCAATTGG |
| EWS2MicroRepeatsFwd | TCTACGACCAATTGAGCGGCCTCTTCCTTCCGATT CTCCAGGAATTCACTGGCCGTCG |
| EWS2MicroRepeatsRevs | CGACGGCCAGTGAATTCCTGGAGAATCGGAAGGA AGAGGCCGCTCAATTGGTCGTAGA |
| EWS4MicroRepeatsFwd | GGTGCTAGAGCTGTCTACGACCAATTGATTCCTTC CTTCCTTCCGATTCTCCAGGAATTCACT |
| EWS4MicroRepeatsRevs | AGTGAATTCCTGGAGAATCGGAAGGAAGGAAGGA ATCAATTGGTCGTAGACAGCTCTAGCACC |
| EWS6MicroRepeatsFwd | GGTGCTAGAGCTGTCTACGATTCCTTCCTTCCTTC CTTCCTTCCGATTCTCCAGGAATTCACTGG |
| EWS6MicroRepeatsRevs | CCAGTGAATTCCTGGAGAATCGGAAGGAAGGAAG GAAGGAAGGAATCGTAGACAGCTCTAGCACC |

**Table S2: $S_{1/2}$ of all the measurements for binding affinities**

| Protein | Substrate | Experiment | $S_{1/2}$ (nM) | # Replicates |
|---|---|---|---|---|
| $FLI1_{85}$ | HA DNA | EMSA | 2.3 ± 0.2 | 3 |
| $FLI1_{85}$ | 2xGGAA DNA | EMSA | 4.3 ± 0.2 | 3 |
| $FLI1_{85}$ | 4xGGAA DNA | EMSA | 5.6 ± 0.9 | 3 |
| $FLI1_{85}$ | 6xGGAA DNA | EMSA | 5.5 ± 0.8 | 3 |
| $FLI1_{85}$ | HA nucleosome | EMSA | 20 ± 7 | 3 |
| $FLI1_{85}$ | 2xGGAA nucleosome | EMSA | 24 ± 2 | 3 |
| $FLI1_{85}$ | 4xGGAA nucleosome | EMSA | 32 ± 2 | 3 |
| $FLI1_{85}$ | 6xGGAA nucleosome | EMSA | 30 ± 2 | 3 |
| $FLI1_{85}$ | HA nucleosome | FRET | 47 ± 6 | 3 |
| $FLI1_{85}$ | 2xGGAA nucleosome | FRET | 92 ± 9 | 3 |
| $FLI1_{85}$ | 4xGGAA nucleosome | FRET | 123 ± 7 | 3 |
| $FLI1_{85}$ | 6xGGAA nucleosome | FRET | 160 ± 10 | 3 |
| $FLI1_{102}$ | HA DNA | EMSA | 2.7 ± 0.2 | 3 |
| $FLI1_{102}$ | 2xGGAA DNA | EMSA | 5.9 ± 0.8 | 3 |
| $FLI1_{102}$ | 4xGGAA DNA | EMSA | 6.7 ± 0.9 | 3 |
| $FLI1_{102}$ | 6xGGAA DNA | EMSA | 5.8 ± 0.5 | 3 |
| $FLI1_{102}$ | HA nucleosome | EMSA | 8 ± 1 | 3 |
| $FLI1_{102}$ | 2xGGAA nucleosome | EMSA | 21 ± 2 | 3 |
| $FLI1_{102}$ | 4xGGAA nucleosome | EMSA | 31 ± 3 | 3 |
| $FLI1_{102}$ | 6xGGAA nucleosome | EMSA | 50 ± 3 | 3 |
| $FLI1_{102}$ | HA nucleosome | FRET | 38 ± 6 | 3 |
| $FLI1_{102}$ | 2xGGAA nucleosome | FRET | 131 ± 9 | 3 |
| $FLI1_{102}$ | 4xGGAA nucleosome | FRET | 153 ± 9 | 3 |
| $FLI1_{102}$ | 6xGGAA nucleosome | FRET | 190 ± 10 | 3 |
| EWSR1::FLI1 | HA DNA | EMSA | 600 ± 400 | 3 |
| EWSR1::FLI1 | 2xGGAA DNA | EMSA | 96 ± 5 | 3 |
| EWSR1::FLI1 | 4xGGAA DNA | EMSA | 94 ± 5 | 3 |
| EWSR1::FLI1 | 6xGGAA DNA | EMSA | 100 ± 10 | 3 |
| EWSR1::FLI1 | HA DNA | FRET | 620 ± 40 | 3 |
| EWSR1::FLI1 | 2xGGAA DNA | FRET | 930 ± 20 | 3 |
| EWSR1::FLI1 | 4xGGAA DNA | FRET | 570 ± 20 | 3 |
| EWSR1::FLI1 | 6xGGAA DNA | FRET | 580 ± 10 | 3 |

**Table S3: FRET Efficiency at 1000 nM**

| Protein | Substrate | FRET Efficiency | # Replicates |
|---|---|---|---|
| $FLI1_{85}$ | HA nucleosome | 0.30 ± 0.01 | 3 |
| $FLI1_{85}$ | 2xGGAA nucleosome | 0.36 ± 0.02 | 3 |
| $FLI1_{85}$ | 4xGGAA nucleosome | 0.31 ± 0.01 | 3 |
| $FLI1_{85}$ | 6xGGAA nucleosome | 0.15 ± 0.01 | 3 |
| $FLI1_{102}$ | HA nucleosome | 0.43 ± 0.01 | 3 |
| $FLI1_{102}$ | 2xGGAA nucleosome | 0.45 ± 0.02 | 3 |
| $FLI1_{102}$ | 4xGGAA nucleosome | 0.40 ± 0.01 | 3 |
| $FLI1_{102}$ | 6xGGAA nucleosome | 0.29 ± 0.01 | 3 |
| EWSR1::FLI1 | HA nucleosome | 0.28 ± 0.01 | 3 |
| EWSR1::FLI1 | 2xGGAA nucleosome | 0.41 ± 0.04 | 3 |
| EWSR1::FLI1 | 4xGGAA nucleosome | 0.20 ± 0.02 | 3 |
| EWSR1::FLI1 | 6xGGAA nucleosome | 0.18 ± 0.01 | 3 |
